# Aerobic capacity and exercise mediate protection against hepatic steatosis via enhanced bile acid metabolism

**DOI:** 10.1101/2024.10.21.619494

**Authors:** Benjamin A. Kugler, Adrianna Maurer, Xiaorong Fu, Edziu Franczak, Nick Ernst, Kevin Schwartze, Julie Allen, Tiangang Li, Peter A. Crawford, Lauren G. Koch, Steven L. Britton, Shawn C. Burgess, John P. Thyfault

## Abstract

High cardiorespiratory fitness and exercise show evidence of altering bile acid (BA) metabolism and are known to protect or treat diet-induced hepatic steatosis, respectively. Here, we tested the hypothesis that high intrinsic aerobic capacity and exercise both increase hepatic BA synthesis measured by the incorporation of ^2^H_2_O. We also leveraged mice with inducible liver-specific deletion of *Cyp7a1* (LCyp7a1KO), which encodes the rate-limiting enzyme for BA synthesis, to test if exercise-induced BA synthesis is critical for exercise to reduce hepatic steatosis. The synthesis of hepatic BA, cholesterol, and *de novo* lipogenesis was measured in rats bred for either high (HCR) vs. low (LCR) aerobic capacity consuming acute and chronic high-fat diets. HCR rats had increased synthesis of cholesterol and certain BA species in the liver compared to LCR rats. We also found that chronic exercise with voluntary wheel running (VWR) (4 weeks) increased newly synthesized BAs of specific species in male C57BL/6J mice compared to sedentary mice. Loss of *Cyp7a1* resulted in fewer new BAs and increased liver triglycerides compared to controls after a 10-week high-fat diet. Additionally, exercise via VWR for 4 weeks effectively reduced hepatic triglycerides in the high-fat diet-fed control male and female mice as expected; however, exercise in LCyp7a1KO mice did not lower liver triglycerides in either sex. These results show that aerobic capacity and exercise increase hepatic BA metabolism, which may be critical for combatting hepatic steatosis.

**Highlights:**

- Rats with intrinsic high aerobic capacity have more significant reductions in *de novo* lipogenesis and increased cholesterol and bile acid synthesis on a high-fat diet compared to rats with low aerobic capacity.
- Chronic exercise increases hepatic bile acid synthesis in mice.
- Loss of *Cyp7a1* blunts the capacity for exercise to increase bile acid synthesis and treat hepatic steatosis in male and female mice fed a high-fat diet.

## 1. INTRODUCTION

Metabolic dysfunction-associated steatotic liver disease (MASLD) is a global epidemic that is associated with metabolic comorbidities [1]. MASLD encompasses a spectrum of liver diseases that begin with the excess accumulation of liver fat (≥5% of liver weight) and can progress to metabolic-associated steatohepatitis (MASH) with inflammation and liver injury. Without intervention, MASLD can lead to irreversible fibrosis (i.e., cirrhosis) and an increased risk of liver cancer (i.e., hepatocellular carcinoma) [2]. Although pharmaceutical treatments for MASLD continue to be evaluated, lifestyle modifications, primarily exercise and dietary changes, remain first-line interventions. In humans, exercise improves aerobic capacity (i.e., cardiorespiratory fitness) while reducing liver triglycerides [3], which is recapitulated in rodent models [4]. Importantly, the effect of exercise to combat hepatic steatosis occurs without weight loss. In addition, lower aerobic capacity independent of body weight has been reported to be associated with MASLD in humans and rodent models [5–8]. However, the mechanisms by which aerobic capacity and exercise prevent and treat hepatic steatosis remain largely unknown.

Elevated fatty acids released from adipose tissue and diet, greater hepatic *de novo* lipogenesis (DNL) from carbohydrates/glucose, and reduced metabolism of fatty acids (fat oxidation, FAO) have all been implicated as causes of MASLD [9]. Utilizing rats bred over several generations for intrinsic aerobic capacity differences, we have shown that high-capacity runners (HCR) have higher hepatic mitochondrial oxidative capacity (i.e., FAO) and are protected from MASLD after exposure to both an acute or chronic high-fat diet (HFD) [6–8]. However, low-capacity runner rats with reduced intrinsic aerobic capacity display lower hepatic oxidative capacity and are highly sensitive to acute and chronic HFD-causing steatosis. Our recent findings demonstrate that HCR rats have elevated gene expression in the cholesterol and bile acid synthesis pathway (i.e., *Hmgcr* and *Cyp7a1*) and increased fecal bile acid loss compared to LCR rats [10; 11]. In addition, aerobic exercise was shown to increase fecal bile acid loss in LDL receptor (*Ldlr*) knockout mice [12]. Further, bile acid sequestrants and the overexpression of cholesterol 7α-hydroxylase (*Cyp7a1*), the rate-limiting enzyme in bile acid synthesis, also increase fecal bile acid loss and protect rodents from steatosis and metabolic derangements of diet-induced obesity [13–15]. However, it is unclear if aerobic capacity and exercise directly upregulate bile acid synthesis and if this is critical for the benefits of exercise in treating MASLD. Thus, we hypothesize that high aerobic capacity and exercise exert their protection against MASLD by promoting bile acid synthesis.

Total bile acid concentration can be quantified by enzymatic assay or by modern liquid chromatography-tandem mass spectrometry (LC-MS/MS), while inference of bile acid synthesis most commonly relies on surrogates of *CYP7A1* enzyme activity, such as 7-hydroxy-4-cholesten-3-one (C4). Here, we used a deuterated water (^2^H_2_O) tracer, which is commonly used to determine the fractional synthesis of lipids, including sterols [16–24], and analogous assumptions have been applied to bile acid synthesis using ^3^H_2_O [25]. Fractional bile acid, cholesterol, and lipid synthesis were quantified in sedentary HCR and LCR rats provided short-term (1 week) and chronic (20 weeks) HFD. Bile acid synthesis was activated in HCR rats in response to HFD. Moreover, these effects were recapitulated by daily exercise (via voluntary wheel running (VWR)) in mice. We further found that inducible liver-specific *Cyp7a1* knockout mice had lower bile acid synthesis and were resistant to the effects of exercise to reduce liver triglycerides, suggesting that exercise-induced bile acid synthesis is critical for the beneficial effects of exercise that treats or protects against steatosis.

## 2. MATERIALS AND METHODS

### 2.1. Ethical approval

All protocols were approved by the Institutional Animal Care and Use Committee at the University of Kansas Medical Center (animal protocol number 2021-2614). All experiments were carried out in accordance with the Guide for the Care and Use of Laboratory Animals published by the National Institutes of Health (NIH Guide, 8th ed., 2011) and adhere to the American Physiological Society’s Guiding Principles in the Care and Use of Vertebrate Animals in Research and Training. Rats were housed in a 12h:12h, dark:light cycle. Both rats and mice were anaesthetized with pentobarbital sodium (100 mg/kg) before a terminal procedure.

### 2.2. High-capacity and low-capacity rat study

The HCR and LCR rat model was developed and characterized at the University of Toledo as previously described [6-8; 26; 27] and shipped to KUMC for the study. At 25–30 weeks of age, animals were singly housed (12:12-h light-dark cycle, 24–26°C). Two different sets of HCR and LCR rats were used for the 1-week (n = 8) and 20-week (n = 10) diet interventions. Only male rats were used in these studies as females do not develop hepatic steatosis on HFD.

During the 1-week study, 64 rats (32 HCRs, 32 LCRs) were acclimatized to the control low-fat diet (LFD; D12110704: 10% kcal fat, 3.5% kcal sucrose, and 3.85 kcal/g, Research Diets, New Brunswick, NJ) for at least 2 weeks before half of each LCR and HCR group (n=16) were transitioned to a high-fat diet (HFD; D12451: 45% kcal fat, 17% kcal sucrose, and 4.73 kcal/g, Research Diets). The other half remained on LFD for 1-week. On the evening before the termination of the experiment, rats were given intraperitoneal ^2^H_2_O injections at a dose of 15uL/g. This dose was estimated to enrich body water to ∼4% ^2^H_2_O. After dosing, rats were subsequently provided 4% ^2^H_2_O drinking water for the remainder of the experiment. Half of the rats from each group in the 1-week study were fasted overnight (∼4pm-8am) (FASTED) while the remaining animals had access to food (FED), allowing us to determine metabolic effects of feeding status across strains. The measurements of food intake, body mass, and body composition (MRI model 900; EchoMRI, Houston, TX) were taken before and after the 1-week intervention. Rats were placed in clean cages just prior to the 1-week diet intervention, and all fecal matter was collected from each cage at the end of the 1-week study. A 20-week study was performed in 40 rats (20 HCRs, 20 LCRs) randomly divided into a HFD and LFD. In the 20-week study all rats had access to food up until tissue collection.

### 2.3. Mouse volunteer wheel running study

Male C57Bl/6J mice (10-12 weeks old; The Jackson Laboratory) were singly housed near thermoneutrality (12:12-h reverse light-dark cycle; ∼30°C) with *ad libitum* access to water and food. Half of the mice were provided with voluntary running wheels (VWR) for 4 weeks, while the other half were maintained in a sedentary condition (n=8 per group). Tissue and serum collection were conducted as described for the rat study, including administration of ^2^H_2_O the night before termination. However, mice were only euthanized in the fed condition.

### 2.4. Liver-specific *Cyp7a1* knockout study

At 10-14 weeks of age, male and female C57BL/6J mice with floxed exons 2-4 of the *Cyp7a1* gene (*Cyp7a1*^fl/fl^, GenePharmatech, Cambridge, MA, T009224) were singly housed at thermoneutrality (∼30C) with *ad libitum* access to a HFD to induce hepatic steatosis. After 4 weeks on the HFD, *Cyp7a1*^fl/fl^ mice were randomly assigned to receive either an intraperitoneal injection of control adeno-associated virus 8 (AAV8)-thyroxin-binding globulin promoter (TBG)-GFP (Control, Ctrl) or AAV8-TBG-Cre leading to liver-specific *Cyp7a1* knockout (LCyp7a1KO). Two weeks post-injection, mice either remained sedentary (SED) or were given access to VWR for daily exercise for 4 weeks to treat hepatic steatosis, resulting in four groups: Ctrl/SED, Ctrl/VWR, LCyp7a1KO/SED, and LCyp7a1KO/VWR (n=6-8 per group in both male and females). Tissue and serum collection were performed as described for the rat study, including administration of ^2^H_2_O the night before termination. However, mice were only euthanized in the fed condition.

### 2.5. Body composition analysis

Body composition and body mass were measured as previously described on the day of tissue collection [8; 28]. Body composition was determined by quantitative magnetic resonance imaging (qMRI) using an EchoMRI-1100 (EchoMRI, TX). Fat-free mass (FFM) was calculated as the difference between body weight and fat mass (FM).

### 2.6. Intestine and fecal total bile acids

The small intestine was frozen and powdered under liquid nitrogen. Rodents received fresh cages 7 days prior to euthanasia, and a total of 7 days of fecal excretion was collected from individual cages. Intestinal tissue (100mg) and feces (100 mg) were weighed, then homogenized using a TissueLyzer II (Qiagen, Germantown, MD) bead homogenizer in 1mL of 100% EtOH. Samples were sealed in parafilm & heated overnight at 50° C, then centrifuged at 1635 ×*g* for 20 minutes. The supernatant was used to measure total bile acid concentration with a commercially available colorimetric kit (DZ042A-KY1/-CAL; Diazyme Laboratories, Inc., Poway, CA). To account for total bile acid content, bile acid concentration was multiplied by total intestinal or fecal weight (from a 1-week collection). Intestinal and fecal bile acid values were corrected for body weight to control for significant differences in body mass.

### 2.7. Fecal energy measurements

Homogenized fecal matter was weighed and pressed into pellets using a Pellet Press (∼600mg) (2811; Parr Instruments, Moline, IL). RO water (2 liters) was weighed out to 2000g ± 0.5g in a calorimetry bucket (A391DD; Parr Instruments, Moline, IL), then placed into a 6100 Compensated Calorimeter (6100EA; Parr Instruments, Moline, IL). Fecal pellets were weighed to 0.0001g and placed into a fuel capsule (43AS; Parr Instruments, Moline, IL). An ignition thread (845DD; Parr Instruments, Moline, IL) was tied to the fuse wire of an Oxygen Combustion Vessel (1108P; Parr Instruments, Moline, IL) before placing the pellet-fuel capsule into the vessel and sealing it. An oxygen supply was connected to the vessel’s inlet valve and then filled to the recommended pressure of 450 psig. After the vessel was pressurized with oxygen, the ignition wires of the calorimeter were connected to the vessel before being placed into the water-filled calorimetry bucket in the calorimeter. The sample ID and mass were entered into the calorimeter prior to starting the system. To account for total energy content, energy concentration was multiplied by total fecal weight (from a 1-week collection) and corrected for body weight.

### 2.8. Serum biological assay

Serum alkaline phosphatase (ALP), aspartate aminotransferase (AST), alanine aminotransferase (ALT), albumin, total protein, blood urea nitrogen (BUN), cholesterol, glucose, and triglyceride measurements were analyzed by a commercial laboratory IDEXX BioAnalytics (North Grafton, MA). Serum β-hydroxybutyrate was determined using a commercially available kit (2440-058; EKF Diagnostics, Boerne, TX). Serum non-esterified fatty acids (NEFAs) were determined using a commercially available kit (NC9517308, -09, -10, -11, -12; FUJIFILM Medical Systems, USA). Serum insulin was determined using a commercially available ELISA kit (80-INSRT-E01; ALPCO, Salem, NH).

### 2.9. Gene expression analysis

RNA was extracted using RNeasy mini-kit following the manufacturer’s instructions (74104; Qiagen, Hilden, DE). Liver gene expression profiles were assessed via bulk RNA-sequencing as previously described [29]. Isolation of polyA RNA and construction of barcoded RNA-seq libraries was performed using TruSeq reagents according to manufacturer’s protocols (Illumina). Quantification of the RNAseq libraries was done using Qubit dsDNA high sensitivity reagents, diluted, denatured, and sequenced using Illumina methodology (HiSeq 2500, 50 bp single reads). Following sequencing and demultiplexing, reads were trimmed for adapters, filtered based on Phred quality score, and aligned to the rat genome using the STAR aligner. Resulting .bam files were imported in Seqmonk for gene level quantification. Differential expression and other analysis including PCA were performed using packages in base R and the limma-voom pipeline. RNA-seq quality metrics including proportion of reads aligning to genic regions were calculated. Pairwise comparisons between HCR and LCR groups within each diet type were performed and differentially expressed genes were identified (p< 0.05, and minimum + 2-fold change). Multiple testing corrections were done using the FDR method. Additional analyses were performed using packages in the R statistical software, ShinyGO app and Gene Set Enrichment Analysis Java application (Broad Institute).

### 2.10. Serum bile acid profiling by LC-MS/MS

Serum bile acid concentrations were quantified by the University of Oklahoma, Laboratory for Molecular Biology and Cytometry Research Metabolomics core (Oklahoma City, OK) using LC-MS methodology as performed previously [30]. 300uL of serum was thawed then vortexed with 600uL of methanol (MeOH) and incubated on ice for 1 hour to precipitate protein. The mixture was centrifuged at 15000 ×g at 4 °C for 20 minutes. Supernatant was transferred to a new Eppendorf tube and dried with Speed-Vacuum. Samples were resuspended in 200 uL of acetonitrile/H_2_O (30:70, v/v) with 0.1% formic acid including 100 ng/ml of G-CDCA-d8 as internal standard, sonicated for 10 minutes in water bath and the supernatant (100 uL) was used for MS analysis.

### 2.11. Tissue bile acid concentration and ^2^H enrichment measured by LC-MS/MS

Briefly, A d9-tauro-chenodeoxycholic acid (d9-TCDCA) internal standard was added to the liver tissues (approximately 30 mg), and the tissue was finely homogenized in 500 µL ice cold MeOH/H_2_O (85:15, v/v) in a 2.0-mL pre-filled Bead Ruptor Tubes (2.8mm ceramic beads, Omni International, Kennesaw, GA, USA). After centrifugation (1635 ×*g* for 10 min) to precipitate the proteins, the supernatant was transferred to a new tube and dried under N_2_. To the dried samples,150 µL of MeOH/H_2_O (50:50, v/v) with 0.1% formic acid was added before MS analysis. Calibration curves were constructed with a fixed amount of d9-TCDCA internal standard. Values for the slope, intercept, and correlation coefficient were obtained by linear-regression analysis of the calibration curves. The area under each analyte peak, relative to the internal standard, was used to calculate the analyte concentrations in liver samples.

LC-MS/MS chromatographic separation of bile acids was performed using a reverse phase C8 column (Phenomenex Luna C8, 150 × 2.0 mm, 3 µm) at a flow rate of 0.2 ml/min. The mobile phase consisted of MeOH/H_2_O (2:98, v/v) with 0.0125% acetic acid (eluent A) and ACN /H_2_O (95:5, v/v) with 0.1% formic acid (eluent B). The gradient proceeded from 25% to 40% B over 12 min and then 40% to 75% B over 12 min. The column was washed with 100% B for 10 min and equilibrated with 25% B for 10 min between injections. Bile acids were detected by an API 3200 triple-quadrupole LC-MS/MS (AB Sciex, MA) operated in negative MRM mode. The ion source parameters were set as follows: curtain gas: 20 psi, ion spray voltage: -4000 V, ion source temperature: 300 °C, and nebulizing and drying gas: 30 and 40 psi. The declustering potential of -120 v, collision Energy of -120 v, entrance potential of -10 v, cell exit potential of -8 v were optimized by infusing each standard solution (1ug/mL). MRM transitions for m0, m1, m2, m3 mass isotopologues of deuterated bile acids, tauro-α-muricholic acid (TαMCA), tauro-β-MCA (TβMCA), taurocholic acid (TCA), tauro-chenodeoxycholic acid (TCDCA), tauro-deoxycholic acid (TDCA) and d9-TCDCA internal standard are summarized in **Supplemental Table 1**. An *m/z* value of 80 (SO ^−^ anion from the taurine moiety) was selected as the common product ion for all the taurine conjugates. Mass to charge (m/z) values of 498.2 (TCDCA and TDCA), 514.2 (TMCA and TCA) and 507.2 (d9-TUDCA) were selected as precursors. Given the existence of isobaric structures in the bile acid pool, we optimized reverse phase LC detection against a mixture of bile acids as reported by Han et al. [31]. Structural isomers, TαMCA, TβMCA and TCA, share the same MRM transitions but were chromatographically separated. TUDCA, TCDCA and TDCA, isomers were also baseline-separated (**Supplemental Fig. S1**).

### 2.12. Liver cholesterol concentration and ^2^H enrichment measured by HR-Orbitrap-GCMS

Approximately 20 mg of tissue was weighed and homogenized with 1 mL of MeOH/DCM (1:1, v/v) in 2.0-mL pre-filled Bead Ruptor Tubes. Tubes were washed twice with 1 mL MeOH/DCM and all solutions were combined. Samples were vortexed and then centrifuged for 5 min at 1635 ×g. A known amount of d7-cholesteroI was added to 2 mg of supernatant and dried under N_2_. Dried extracts were saponified with 1 mL 0.5 M KOH in MeOH at 80°C for 1 h. Lipids were extracted with DCM/water before evaporation to dryness. The dried lipid extract was derivatized by incubation at 75°C for 1 h addition with 100 µL acetyl chloride. The sample was evaporated to dryness under N_2_ and was reconstituted in 100 µL iso-octane for analysis by GCMS.

The ^2^H-enrichment of cholesterol (m0, m1, m2, m3 isotopologues of deuterated cholesterol) was determined using a Q Exactive GC-orbitrap MS (Thermo Scientific). 1 μL of sample was injected onto a HP-5ms capillary column (60m×0.32mm i.d., 0.25µm film thickness) in split mode. Helium gas flow rate was set to 13.5 min of 1 mL/min for the initial injection, followed by 0.4mL/min for 5 min before returning to 1 mL/min. The GC injector temperature was set at 250°C and the transfer line was held at 290°C. The column temperature was set to 200°C for 1 min and increased by 20°C/min before reaching 320°C over 16 min. Samples were analyzed at 70 eV in EI mode by targeted selected ion monitoring (t-SIM) at 240,000 mass resolution (FWHM, m/z 200). Tuning and calibration of the mass spectrometer was performed using perfluorotributylamine (FC-43) to achieve a mass accuracy of <0.5 ppm. The quadrupole was set to pass ions between m/z 246.24 and 252.24. The Orbitrap automatic gain control (AGC) target was set to 5e^4^ with a maximum injection time of 54 ms. Cholesterol concentration was calculated from the area ratio of the peaks corresponding to cholesterol (m/z 247.242) and the D7-cholesterol internal standard (m/z 254.286) with full scan mass ranges 240-260 m/z. Extraction of individual high-resolution m/z values representing each isotopomer ion was done using TraceFinder 4.1 (Thermo Scientific) with 4 ppm mass tolerance.

### 2.13. Triglyceride palmitate concentration and ^2^H enrichment measured by HR-Orbitrap-GCMS

Liver palmitate was measured as previously reported [18] and followed the same sample preparation as described for cholesterol analysis, except the dried lipid extract was resuspended in 50 µL of 1% triethylamine/acetone and reacted with 50 µL of 1% Pentafluorobenzyl bromide/acetone for 30 minutes at room temperature. To this solution, 1 mL of iso-octane was added before MS analysis. The ^2^H-enrichment of palmitate was determined using HR-Orbitrap-GCMS as previously described [18].

### 2.14. Body water enrichment measured by HR-Orbitrap-GCMS

Plasma samples dissolved in acetone under alkaline conditions directly in the autosampler vial as previously reported [18]. In brief, 5 μl of plasma sample, 2 μl of 10 M sodium hydroxide, and 5 μl of acetone were added to a threaded GC vial. Samples were incubated overnight at room temperature prior to analysis. Calibration standards of known ^2^H-mol fraction excess were prepared by mixing weighed samples of natural abundance and of 99.9% ^2^H_2_O. Negative chemical ionization mode (NCI) was used with t-SIM acquisition (m/z 55.5-60.5) and 60,000 mass resolution (FWHM, m/z 200) on the same HR-Orbitrap-GCMS instrument as described previously [18].

### 2.15. Fractional synthesis of palmitate, cholesterol, and bile acids

The fractional synthesis of palmitate, cholesterol and bile acids were calculated as:

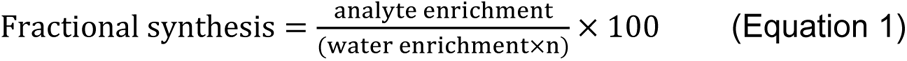

Palmitate, cholesterol, and bile acid analyte ^2^H enrichment was determined from mass isotopomers m1 (^2^H_1_), m2 (^2^H_2_), and m3 (^2^H_3_), as described above, and correction for naturally abundant isotopes was made using the MID of a biological sample (collected without ^2^H_2_O administration) and a matrix correction algorithm. Analyte ^2^H enrichment = ^2^H_1_ + (^2^H_2_ x 2) + (^2^H_3_ x 3). *N* is the number of deuterium exchangeable hydrogens in each analyte and can be experimentally determined from the binomial distribution of their MIDs [21; 24; 32]. Palmitate was previously found to have n=22 [18]. The partial cholesterol fragment (m/z 247) was determined to have n=20, which is proportionally similar to the full cholesterol ion previously reported [21]. Assignment for TaMCA n=14, TbMCA n=10, TCA n=14, TCDCA n=18, TDCA n=10 were made from their MIDs based on the assumption of normal binomial distributions.

### 2.16. Measurement of liver triglycerides

Intrahepatic triglycerides (TAG) as previously described [6; 26]. In brief, hepatic lipids were extracted by adding 1 mL of 1:2 vol/vol methanol-chloroform to powdered liver tissue (∼30 mg). The mixture was homogenized and rotated overnight at 4°C. Then, 1 mL of 4 mM MgCl₂ was added, and the sample was centrifuged for 1 hour at 1,000 g at 4°C. The organic phase was collected, evaporated, and reconstituted in a 3:2 vol/vol butanol-Triton X-114 mix. Hepatic TAGs were measured using a commercially available kit (Sigma, TR0100-1KT). Data were then normalized to liver weight.

### 2.17. Statistics

Measurements at 1-week and 20-week in the HCR and LCR rats were analyzed independently. Anthropometrics and energy intake are only reported for the FED groups from the 1-week study and were analyzed using 2-way ANOVA (strain X diet) followed by Tukey’s multiple comparisons test. Intestinal, liver, and serum bile acids, serum metabolites, and DNL and cholesterol synthesis were analyzed using 3-way ANOVA (strain X diet X fed state) followed by Tukey’s multiple comparisons test. All fecal measurements were corrected for body weight due to significant differences in body mass between HCR and LCR rats. FED and FASTED fecal measurements during the 1-week study were pooled together (animals were only fasted 1 night prior to sacrifice while feces was collected over 7 days) according to strain and diet then analyzed using 2-way ANOVA (strain X diet) followed by Tukey’s multiple comparisons test. All 20-week measurements, except bile acid synthesis, were analyzed using 2-way ANOVA (strain X diet) followed by Tukey’s multiple comparisons test. In wild-type mice studies, comparisons of bile acid synthesis between VWR vs. sedentary were made via unpaired T-test. Bile acid synthesis measurements were analyzed using unpaired t-test. LCyp7a1KO was analyzed within sex utilizing 2-way ANOVA (Genotype X VWR). Statistical analyses were performed in Prism 10 (GraphPad Software, San Diego, CA).

## 3. RESULTS

### 3.1. HCR rats display less weight gain and changes in circulating lipids on a high-fat diet

As expected, body mass and percent fat mass were greater, while the percent lean mass was reduced in LCR strain at the end of the 1-week than HCR counterparts (Main effect of strain, P<0.05, **Supplemental Table S2**). One week of diet significantly increased body mass, which was influenced by increases in fat mass (Main effect of Diet, P<0.05). However, a diet and strain interaction revealed this was driven by the LCR fed a HFD as they had a significantly greater increase in fat mass (P<0.05), which was not observed in the HCR rats fed HFD. This effect was influenced by increased energy intake from the diet (Main effect of diet, P<0.05) by LCR rats on the HFD (P<0.05), which was not observed in HCR rats.

After 20 weeks of HFD, body mass, and percent fat mass were greater, while percent lean mass was reduced in the LCR rats compared to HCR rats (Main effect of strain, P<0.05, **Supplemental Table S3**). Twenty weeks of a HFD increased body mass and percent fat mass and reduced percent lean mass regardless of strain, which was influenced by increased energy intake (Main effect of diet, P<0.05). However, strain and diet interactions revealed that changes in body mass, fat mass, and energy intake were driven by the significant increase in the LCR rats fed a HFD compared to LCR rats fed a LFD (P<0.05).

Serum metabolic data for 1-week and 20-week diets are shown in **Supplemental Table S4** and **Table S5**. Nutritional state (i.e., Fed vs Fasted) affected all variables except serum NEFAs in the 1-week HFD condition. Surprisingly, ALP, AST, and ALT were significantly lower in LCR rats than matched HCR rats (Main effect of strain, P<0.05). Serum cholesterol, triglycerides, and NEFA were higher in LCR compared to HCR rats (Main effect of strain, P<0.05). Serum insulin was generally lower in LCR rats than in HCR counterparts (Main effect of strain, P<0.05).

Differences in ALP, AST, and ALT between strains disappeared after the 20-week HFD. HCRs had lower serum cholesterol than LCR counterparts after 20 weeks of the diet (Main effect of strain, P<0.05, **Supplemental Table S5**). Serum β-hydroxybutyrate and NEFAs were increased in both strains fed a HFD (Main effect of diet, P<0.05). There was an interaction between strain and diet for serum triglycerides, as serum triglycerides were elevated in HCR rats on an LFD compared to LCR rats fed an LFD (P<0.05). However, LCR rats fed a HFD had a significant increase in serum triglycerides compared to LCR fed an LFD (P<0.05).

### 3.2. HCR and LCR rats display different serum and liver bile acid levels and composition

One week after diet intervention, serum total bile acids were significantly lower in HCR rats compared to LCR rats, regardless of diet (Main effect of strain, P<0.05, **Fig. 1A and Supplemental Table S6**). Due to variations in the serum bile acid pool size, conjugated and unconjugated bile acids were analyzed as a percentage of the total serum bile acid pool. Glycine-conjugated bile acids were higher in LCR rats than HCR rats (Main effect of strain, P<0.05, **Fig. 1A and Supplemental Table S6**). Fasting led to a higher proportion of glycine-conjugated bile acids in the LCR rats (Main effect of fasting, P<0.05, **Fig. 1A and Supplemental Table S6**). Interestingly, the 12α-hydroxylated to non-12α-hydroxylated bile acid ratio was elevated during fasting conditions (Main effect of fasting, P<0.05, **Supplemental Table S6**), but this response was largely driven by a change induced by fasting in the LCR, suggesting an increase in classical or a decrease in alternative bile acid synthesis due to fasting in LCR rats.

**Figure 1.**
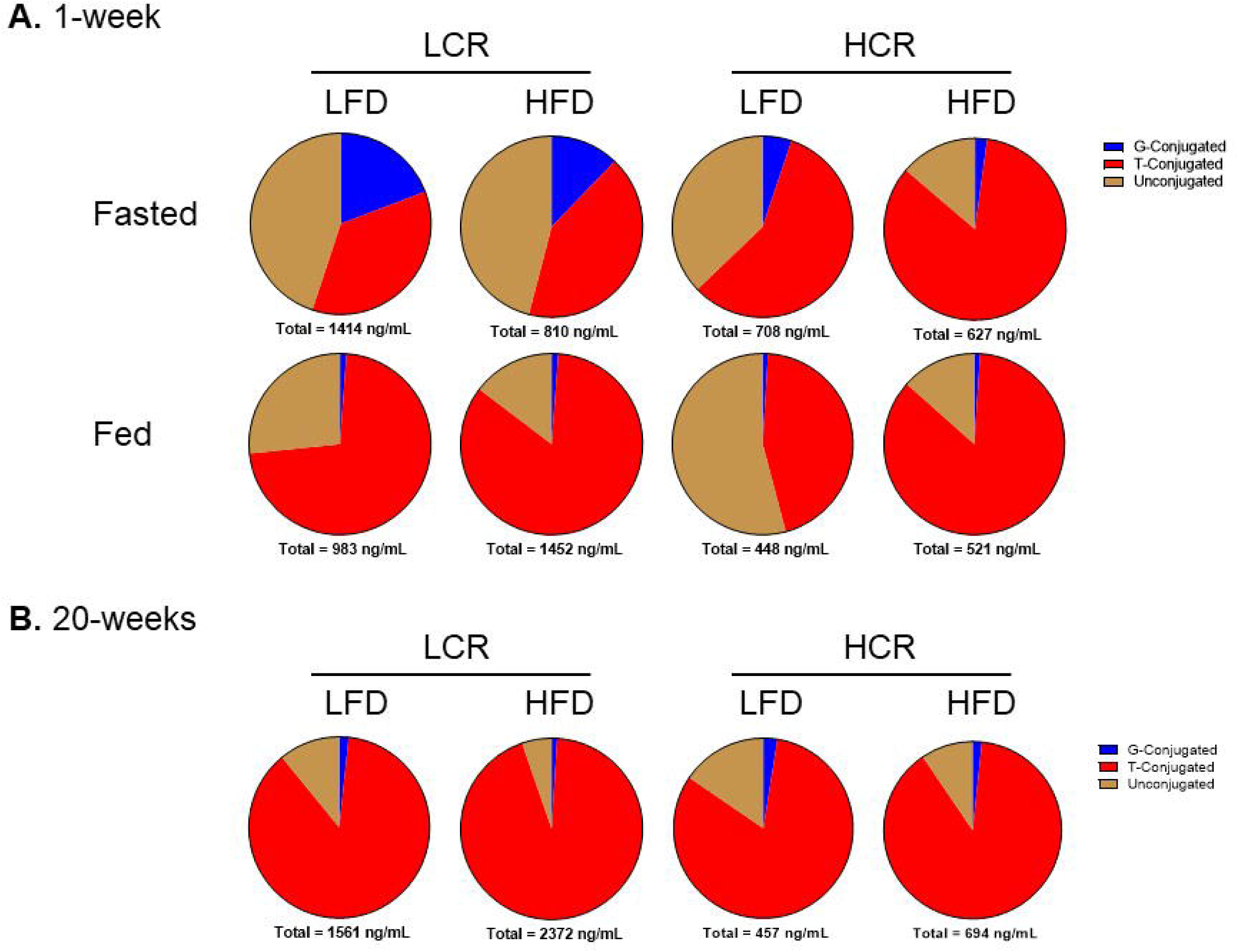
Serum Bile Acid Composition. **A.** Serum bile acid composition and total bile acids from rats during a 1-week study (n=8). **B.** Serum bile acid composition and total bile acids from rats during a 20-week study (n=10).

Liver bile acid measurements focused specifically on taurine-conjugated bile acids because they comprise the largest proportion of the bile acid pool in rodents. Total liver bile acid concentration was higher in the LCR rats after the 1-week diet intervention (Main effect of strain, P<0.05, **Table 1**). Specifically, T-αMCA and T-CA concentrations were greater in LCR than HCR counterparts (Main effect of strain, P<0.05, **Table 1**). However, fasting increased liver bile acid content, particularly T-CA and T-DCA, in both strains (Main effect of fasting, P<0.05, **Table 1**).

**Table 1.** Liver bile acid concentrations from HCR/LCR rats on a LFD or HFD for 1-week.

| (ug/g of liver) | LCR |  |  |  | HCR |  |  |  | P-value |  |  |  |  |
| --- | --- | --- | --- | --- | --- | --- | --- | --- | --- | --- | --- | --- | --- |
|  | FASTED |  | FED |  | FASTED |  | FED |  | Strain | Fed | Diet | Strain x Fed | Strain x Diet |
|  | LFD | HFD | LFD | HFD | LFD | HFD | LFD | HFD |  |  |  |  |  |
| T-αMCA | 8.2±1.5 | 7.1±1.4 | 5.4±0.6 | 8.3±1.51 | 5.0±0.6 | 3.7±0.9 | 5.9±1.4 | 4.1±0.8 | **0.003** | 0.924 | 0.721 | 0.404 | 0.135 |
| T-βMCA | 13.3±3.2 | 19.7±9.1 | 32.3±4.5 | 35.0±5.1 | 25.5±3.5 | 22.2±2.8 | 19.7±4.0 | 27.1±6.3 | 0.696 | **0.031** | 0.382 | **0.023** | 0.738 |
| T-CA | 112.9±16.7 | 114.7±22.5 | 48.6±7.2<sup>‡‡</sup> | 71.9±10.2 | 71.4±9.8 | 82.3±8.6 | 29.5±6.1 | 51.6±10.4 | **0.002** | **<0.001** | 0.091 | 0.311 | 0.814 |
| T-CDCA | 10.9±2.6 | 9.7±1.6 | 5.6±0.7 | 10.1±1.6 | 7.3±1.7 | 6.8±1.7 | 8.4±1.6 | 6.0±0.8 | 0.077 | 0.291 | 0.914 | 0.251 | 0.154 |
| T-DCA | 11.5±4.7 | 9.3±1.9 | 2.2±0.4<sup>‡</sup> | 2.3±0.6 | 10.4±3.1 | 10.2±0.8 | 3.2±0.7 | 2.8±0.5 | 0.830 | **<0.001** | 0.615 | 0.734 | 0.781 |
| **TOTAL** | 156.8±17.6 | 160.6±34.9 | 94.1±11.6 | 127.7±16.9 | 119.5±13.4 | 125.2±12.2 | 66.6±10.0 | 91.5±17.5 | **0.009** | **<0.001** | 0.184 | 0.858 | 0.892 |

Values are means ± SEM (n=6-8). Significance was determined by 3-way ANOVA (strain X diet X fed state) followed by Tukey's multiple comparisons test; ‡ indicates effect of fed state within strain and diet (‡p<0.05, ‡‡p<0.01, ‡‡‡p<0.001).

After 20 weeks of a HFD, serum bile acid concentration increased in LCR rats but not in HCR (Main effect of strain, P<0.05, **Fig. 1B and Supplemental Table S7**). This increase was driven by elevated T-βMCA, T-CA, T-DCA, and T-UDCA in LCR (Main effect of strain, P<0.05, **Supplemental Table S7**). Again, the 12α-hydroxylated to non-12α-hydroxylated bile acid ratio was higher in LCR rats than in HCR counterparts (Main effect of strain, P<0.05, **Supplemental Table S7**); however, regardless of strain, HFD also increased the 12α-hydroxy/non-12α-hydroxy ratio (Main effect of diet, P<0.05, **Supplemental Table S7**). Total glycine- and taurine-conjugated bile acids were elevated in LCR rats compared to HCR rats (Main effect of strain, P<0.05, **Supplemental Table S7**). Despite these differences, the serum bile acid percent composition was not significantly different between HCR and LCR rats after chronic HFD **(Fig. 1B**). Similar to serum bile acids, liver bile acids were elevated in LCR rats fed an HFD compared to HCR counterparts (Main effect of strain, P<0.05, **Table 2**) an effect driven by increased T-βMCA and T-CA in LCR (Main effect of strain, P<0.05, **Table 2**).

**Table 2.** Liver bile acid concentrations from HCR/LCR rats on only a HFD for 20-weeks.

| (ug/g of liver) | LCR | HCR | P-value |
| --- | --- | --- | --- |
| T-αMCA | 5.06 ± 0.7 | 3.14 ± 0.80 | 0.090 |
| T-βMCA | 42.24 ± 8.75 | 17.44 ± 4.90 ^ | **0.024** |
| T-CA | 118.30 ± 14.52 | 52.79 ± 9.49 ^^ | **0.001** |
| T-CDCA | 4.08 ± 0.69 | 3.80 ± 0.74 | 0.781 |
| T-DCA | 4.12 ± 0.86 | 3.56 ± 0.48 | 0.573 |
| **TOTAL** | 173.80 ± 23.90 | 80.73 ± 15.80 ^^ | **0.005** |

Values are means ± SEM (n=10). Significance was determined by Student's unpaired t-test; ^indicates effect of strain (^p<0.05, ^^p<0.01, ^^^p<0.001).

### 3.3. HCR rats have increased fecal bile acids and energy loss

After correcting for body weight, intestinal bile acids were not different between LCR and HCR rats in either the 1-week or 20-week study (**Fig. 2A and B**). However, fecal bile acid content was significantly higher in HCR rats compared to LCR rats in both diet conditions (Main effect of strain, P<0.05, **Fig. 2C and D**). HCR also had higher fecal energy loss in both diet conditions than LCR (Main effect of strain, P<0.05, **Fig. 2E and F**).

**Figure 2.**
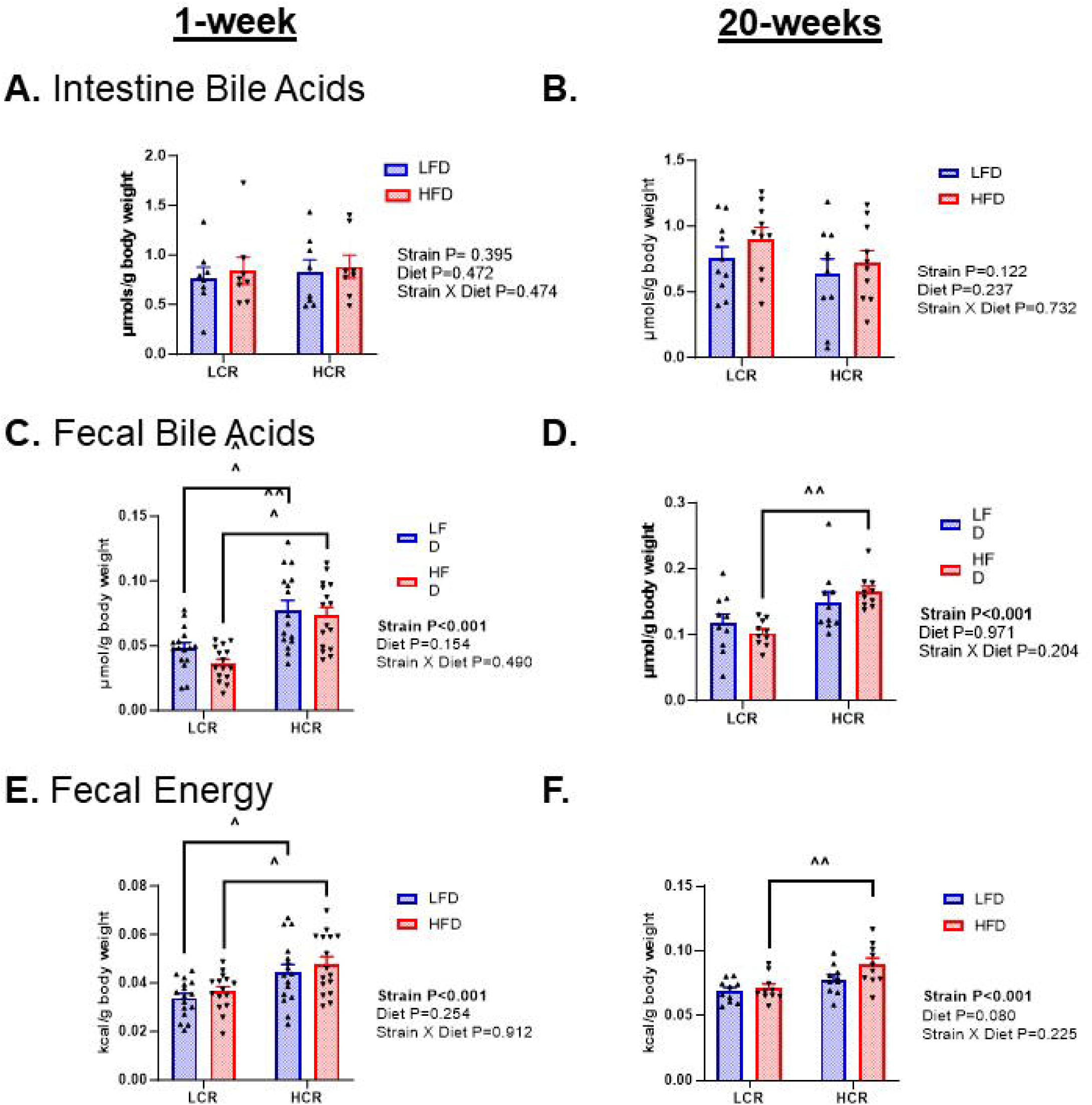
Intestinal and Fecal Bile Acid Content and Fecal Energy Loss. **A.** Intestinal bile acid measurements from rats during the 1-week study (n=8). **B.** Intestinal bile acid measurements from rats during the 20-week study (n=10). **C.** Fecal bile acid content from rats during the 1-week study (n=16). **D.** Fecal bile acid content from rats during the 20-week study (n=10). **E.** Fecal energy loss from rats during the 1-week study (n=16). **F.** Fecal energy loss from rats during the 20-week study (n=10). Data represented as means ± SEM. ^indicates effect of strain within diet (^p<0.05, ^^p<0.01, ^^^p<0.001).

### 3.4. HCR rats have greater cholesterol and bile acid synthesis

Consistent with our previous findings in mice [33], a 1-week HFD suppressed *de novo* lipogenesis (DNL) compared to a LFD in both strains (Main effect of diet, P<0.05, **Fig. 3A**). Interestingly, HCR rats had a more robust reduction in DNL than the LCR after the 1-week HFD (P<0.05, **Fig. 3A**). Hepatic cholesterol synthesis was higher in HCR rats compared to LCR rats (Main effect of strain, P<0.05, **Fig. 3B**), which was partially driven by higher cholesterol synthesis during fasting (Main effect of fasting, P<0.05, **Fig. 3B**). DNL and cholesterol synthesis were not measured in the 20-week HFD condition.

**Figure 3.**
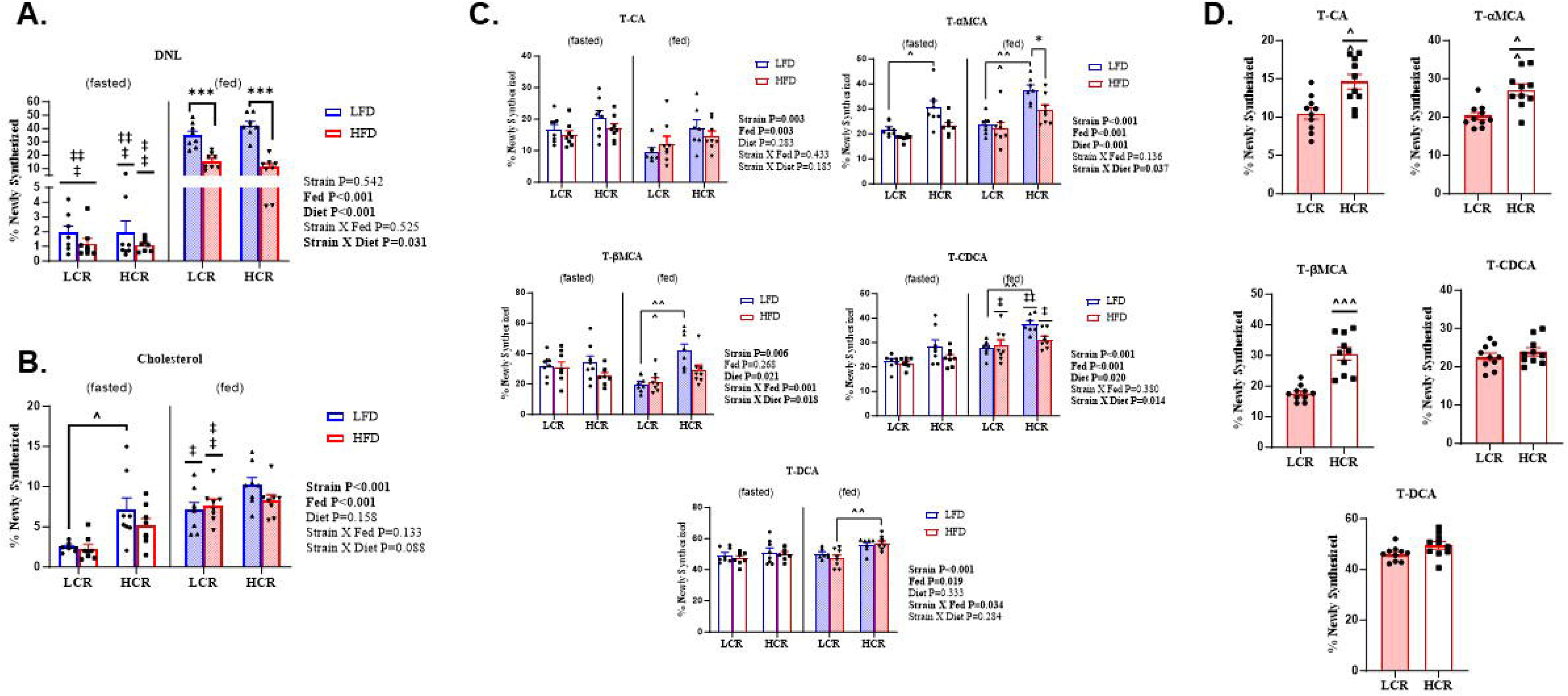
DNL, Cholesterol Synthesis, and Bile Acid synthesis. **A.** DNL as measured by ^2^H incorporation into % newly synthesized hepatic palmitate from rats during the 1-week study (n=8). **B.** Cholesterol synthesis as measured by ^2^H incorporation into % newly synthesized hepatic cholesterol from rats during a 1-week study (n=8). **C.** Bile acid synthesis as measured by ^2^H incorporation into % newly synthesized T-αMCA, T-βMCA, T-CA, T-CDCA, and T-DCA from rats during a 1-week study (n=6-8). **D.** Bile acid synthesis as measured by ^2^H incorporation into % newly synthesized T-αMCA, T-βMCA, T-CA, T-CDCA, and T-DCA from rats during a 20-week study (n=10). Data represented as means ± SEM. *indicates effect of diet within strain and fed state (*p<0.05, **p<0.01, ***p<0.001); ^indicates effect of strain within diet and fed state (^p<0.05, ^^p<0.01, ^^^p<0.001); ‡indicates effect of fed state within strain and diet (‡p<0.05, ‡‡p<0.01, ‡‡‡p<0.001).

Newly synthesized bile acids T-CA, T-αMCA, T-βMCA, T-CDCA, and T-DCA were higher in HCR rats compared to LCR counterparts after the 1-week HFD (Main effect of strain, P<0.05, **Fig. 3C**). Overnight fasting reduced the synthesis of the majority of bile acids except for T-CA, which was increased (Main effect of fasting, P<0.05, **Fig. 3C**). The 1-week HFD reduced bile acid synthesis in both strains and in both fasted/fed conditions (Main effect of diet, P<0.05, **Fig. 3C**). Following the 20-week HFD, the percentage of newly synthesized bile acids was higher in HCR rats than in LCR rats; as primary bile acids T-αMCA, T-βMCA, and T-CA were statistically significant (Main effect of strain, P<0.05, **Fig. 3D**). These data show that elevated bile acid synthesis in HCR over the LCR is maintained over the course of a long term HFD.

### 3.5. Aerobic capacity regulates hepatic bile acid gene expression

We have previously reported that HCR displays upregulated transcription of cholesterol and bile acid synthesis pathways in the liver than LCR [10; 11]. Similarly, HMG-CoA reductase gene (*Hmgcr*) expression was higher in HCR rats (Main effect of strain, P<0.05, **Fig. 4A**) as was gene expression for the rate-limiting enzyme of bile acid synthesis, *Cyp7a1*, and the alternative pathway, *Cyp27a1* (Main effect of strain, P<0.05, **Fig. 4B and D**). Hepatic *Cyp8b1* expression was not different between strains on the 1-week HFD study when fasted, but in the fed condition, HCR displayed higher expression than LCR (Main effect of diet, P<0.05, **Fig. 4C**). Hepatic *Cyp7b1,* which his downstream of *Cyp27a1* was lower in HCR than LCR across all conditions (Main effect of strain, P<0.05, **Fig. 4E)** as was *Baat* expression, an enzyme that regulates conjugation of bile acids (Main effect of strain, P<0.05, **Fig. 4F**).

**Figure 4.**
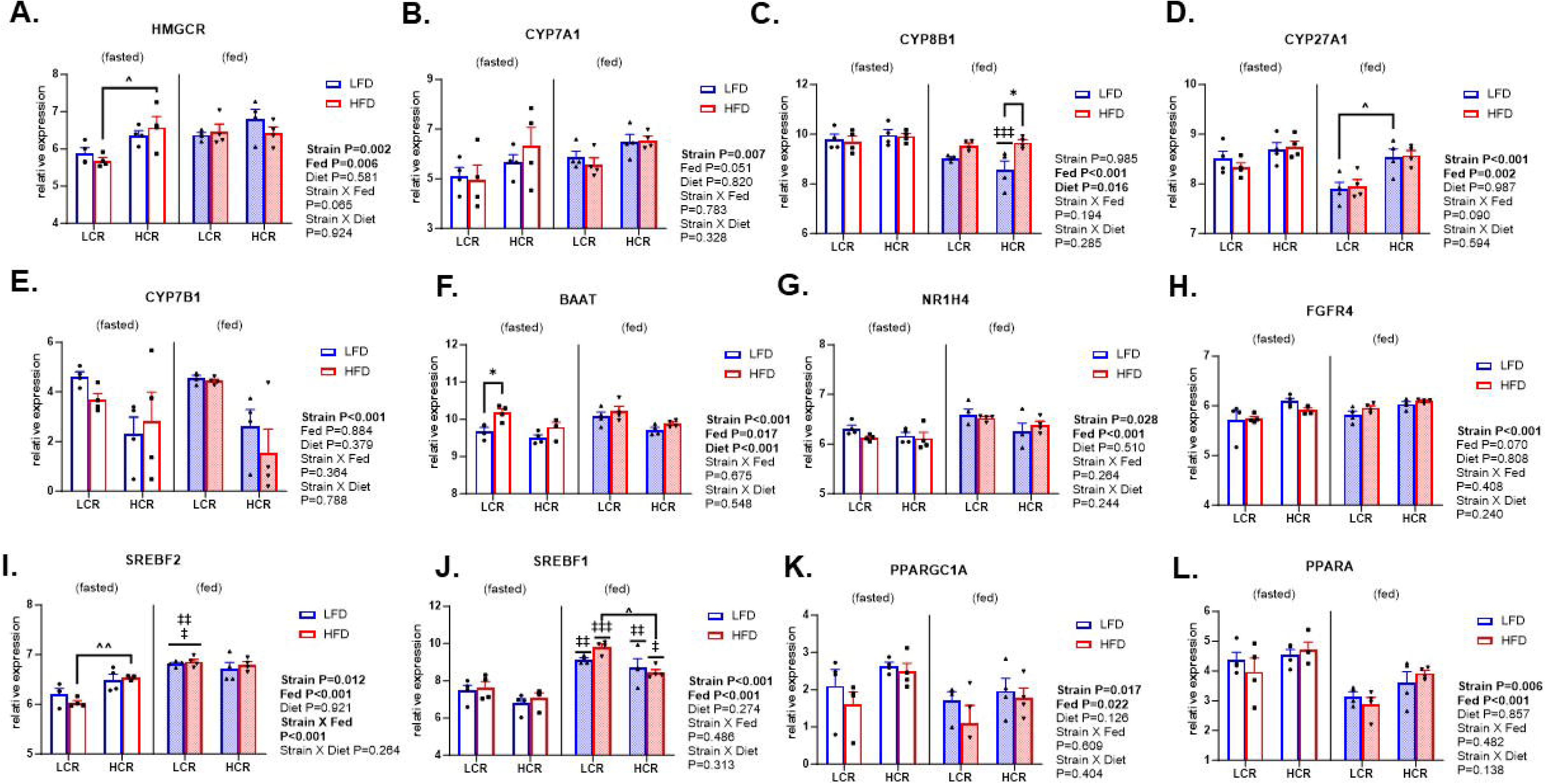
Cholesterol and Bile Acid Synthesis Gene Expression in HCR and LCR rats during a 1-week study. **A.** Gene expression for the cholesterol synthesis protein, HMG-CoA reductase (HMGCR). **B.** Gene expression for the rate-limiting protein in bile acid synthesis, Cyp7a1. **C.** Gene expression for a mitochondrial protein involved in the bile acid synthetic pathway, Cyp27a1. **D.** Gene expression for the protein responsible for determining bile acid pool composition, Cyp8b1. **E.** Gene expression for a protein in the alternative bile acid synthetic pathway, Cyp7b1. **F.** Gene expression for the Bile Acid-CoA:Amino Acid N-Acyltransferase (BAT) enzyme which controls the conjugation of bile acids to an amino acid synthesis (BAAT). **G.** Gene expression for the hepatic nuclear receptor involved in redundant feedback regulation of bile acids, FXR (NR1H4). **H.** Gene expression for a hepatic receptor involved in bile acid feedback from the intestines, FGFR4. **I.** Gene expression for a transcription factor that promotes cholesterol synthesis, SREBP-2. **J.** Gene expression for a mitochondrial protein involved in the bile acid synthetic pathway, SREBF1. **K.** Gene expression for the transcriptional co-activator peroxisome gamma co-activator 1 alpha (PGC1α), a master regulator of mitochondrial biogenesis and genes involved in energy metabolism (PGC1α). **L.** Gene expression for a transcription factor that helps regulate fatty acid oxidation in the liver, PPARα (PPARα). Data represented as normalized gene expression values with units as log-transformed counts per million (means ± SEM; n=4). *Indicates effect of diet within strain and fed state (*p<0.05, **p<0.01, ***p<0.001); ^indicates effect of strain within diet and fed state (^p<0.05, ^^p<0.01, ^^^p<0.001); ‡indicates effect of fed state within strain and diet (‡p<0.05, ‡‡p<0.01, ‡‡‡p<0.001).

Bile acid synthesis is regulated by a negative feedback loop in which bile acids returning to the liver activate the nuclear receptor FXR to suppress *Cyp7a1* expression. Liver FXR (encoded by the *Nr1h4* gene) was lower in HCR rats compared to LCR rats (Main effect of strain, P<0.05, **Fig. 4G**). In contrast, another regulator of bile acid and cholesterol synthesis, *Fgfr4*, was higher in HCR than LCR regardless of diet (Main effect of strain, P<0.05, **Fig. 4H**). Consistent with the differences found for cholesterol synthesis between strains, *Srebp-2* (encoded by the *Srebf-2* gene) expression was consistently higher in HCR vs. LCR (Main effect of strain, P<0.05, **Fig. 4I**). However, a strain and fasting interaction revealed that this difference was driven by lower *Srebp-2* gene expression in fasting LCR rats (P<0.05). Liver *Srebf-1* expression, which encodes for *Srebp-1*, was induced in both strains in the fed state (Main effect of fasting, P<0.05, **Fig. 4J**) and remained higher in LCR across all diets/conditions (Main effect of strain, P<0.05, **Fig. 4J**). As expected, due to their known higher mitochondrial oxidative capacity, HCR rats had higher hepatic gene expression of the transcriptional co-activator peroxisome gamma co-activator 1 alpha (*Pgc1α*) and peroxisome proliferator-activated receptor alpha (*Pparα*), regardless of diet or fasting condition (Main effect of strain, P<0.05, **Fig. 4K and L**).

### 3.6. Exercise via VWR increases bile acid synthesis in mice

Because exercise can increase aerobic capacity, we next examined whether chronic exercise increases hepatic bile acid metabolism in male mice and recapitulates the contrasting responses in HCR vs. LCR rats. After 4 weeks, VWR reduced body weight and fat mass while increasing lean body mass and energy intake (P<0.05, **Supplemental Table S8**). Remarkably, VWR increased bile acid synthesis by elevating the synthesis of primary bile acids T-CA, T-αMCA, T-βMCA, and T-CDCA, and secondary bile acid T-DCA, which correlated with increased T-CA (P<0.05, **Fig. 5A-E**) compared to sedentary control mice. These data confirm the induction of bile acid synthesis in response to exercise training.

**Figure 5.**
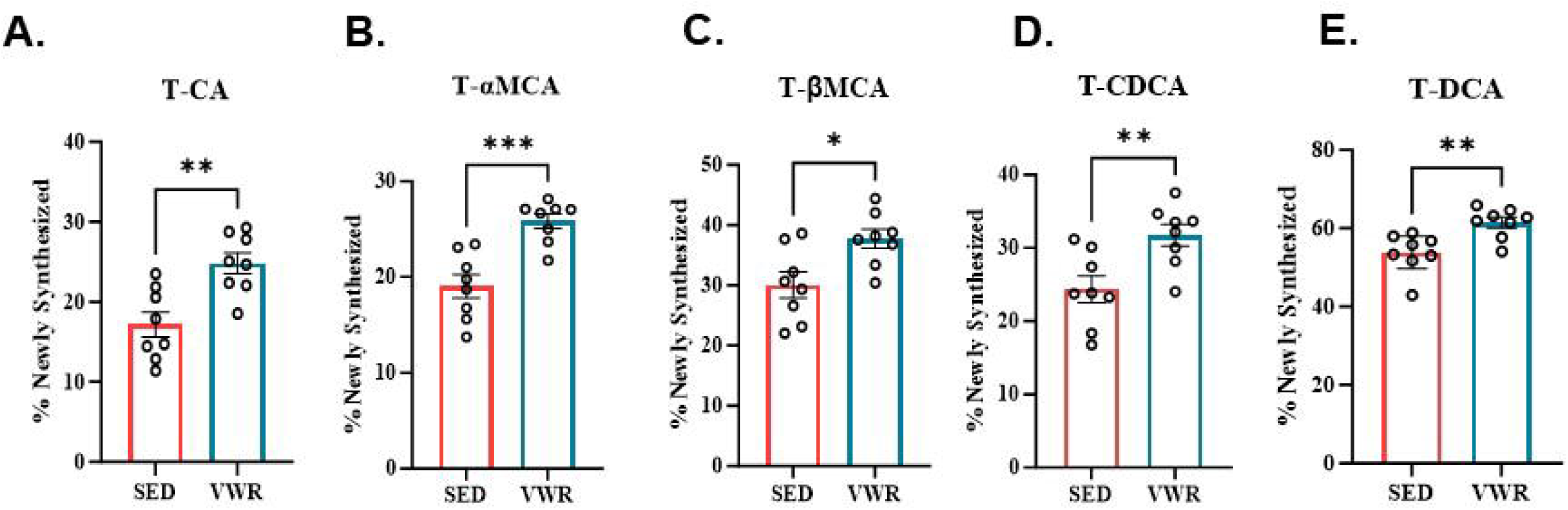
Bile Acid Synthesis Measures in VWR Mice. Data shows bile acid synthesis as measured by ^2^H incorporation into % newly synthesized **A.** T-CA. **B.** T-αMCA. **C.** T-βMCA. **D.** T-CDCA and **E.** T-DCA. Measurements from mice (n=8) on a HFD (Control) that either remained sedentary (SED) or were given running wheels (VWR) for 4 weeks. Data represented as means ± SEM. *indicates effect of diet within running or sedentary condition (*p<0.05, **p<0.01, ***p<0.001); ^indicates effect of running or sedentary condition within diet (^p<0.05, ^^p<0.01, ^^^p<0.001).

### 3.7. *Cyp7a1* mediated bile acid synthesis is critical for exercise to treat steatosis

We and others have shown that exercise protects and treats HFD-induced hepatic steatosis in mice [34]. In the current and previous studies, we reported that *Cyp7a1* gene expression is upregulated in HCR rats on a HFD and in exercising mice, indicating that *Cyp7a1* is a critical factor in the ability of exercise to prevent hepatic steatosis [10; 11]. We also found that exercise in rats and mice increases hepatic expression of genes regulating bile acid and cholesterol synthesis (*Acly*, *Cyp7a1*, and *Hmgcr*), suggesting that bile acid synthesis is upregulated by exercise [11]. To investigate these effects further, we developed an inducible liver-specific *Cyp7a1* knockout mouse model in which *Cyp7a1* expression was knocked out in the liver before exercise. The LCyp7a1KO had reduced *Cyp7a1* gene expression, confirming the liver-specific knockout of *Cyp7a1* (Main effect of LCyp7a1KO, P<0.05, **Fig 6A**). There were no differences in body weight or body composition in male mice during the 4 weeks of exercise regardless of genotype (**Supplemental Table S9**), but exercise increased daily energy intake as usual (Main effect of VWR, P<0.05, **Supplemental Table S9**). Similarly, female mice showed no difference in body weight regardless of exercise or genotype (**Supplemental Table S10**), however, exercise reduced fat mass while increasing energy intake (Main effect of VWR, P<0.05, **Supplemental Table S10**). Overall, liver-specific *Cyp7a1* knockout did not alter weight gain or body composition. Liver triglycerides were significantly elevated in LCyp7a1KO mice compared to control, regardless of sex or exercise (Main effect of LCyp7a1KO, P<0.05, **Fig 6B and C**). While liver triglycerides tended to be reduced by exercise in both male (-45.5%) and female (-52.8%) control groups, this effect was not observed in LCyp7a1KO mice. Liver content of the bile acids T-CA, T-αMCA, T-CDCA, and T-DCA were all significantly reduced in LCyp7a1KO mice of both sexes compared to controls (Main effect of LCyp7a1KO, P<0.05, **Supplemental Tables S11 and S12**). Moreover, the total bile acid content in the liver, gallbladder, intestines, and feces was significantly lower in LCyp7a1KO mice (Main effect of LCyp7a1KO, P<0.05, **Supplemental Fig 2A-D**).

**Figure 6.**
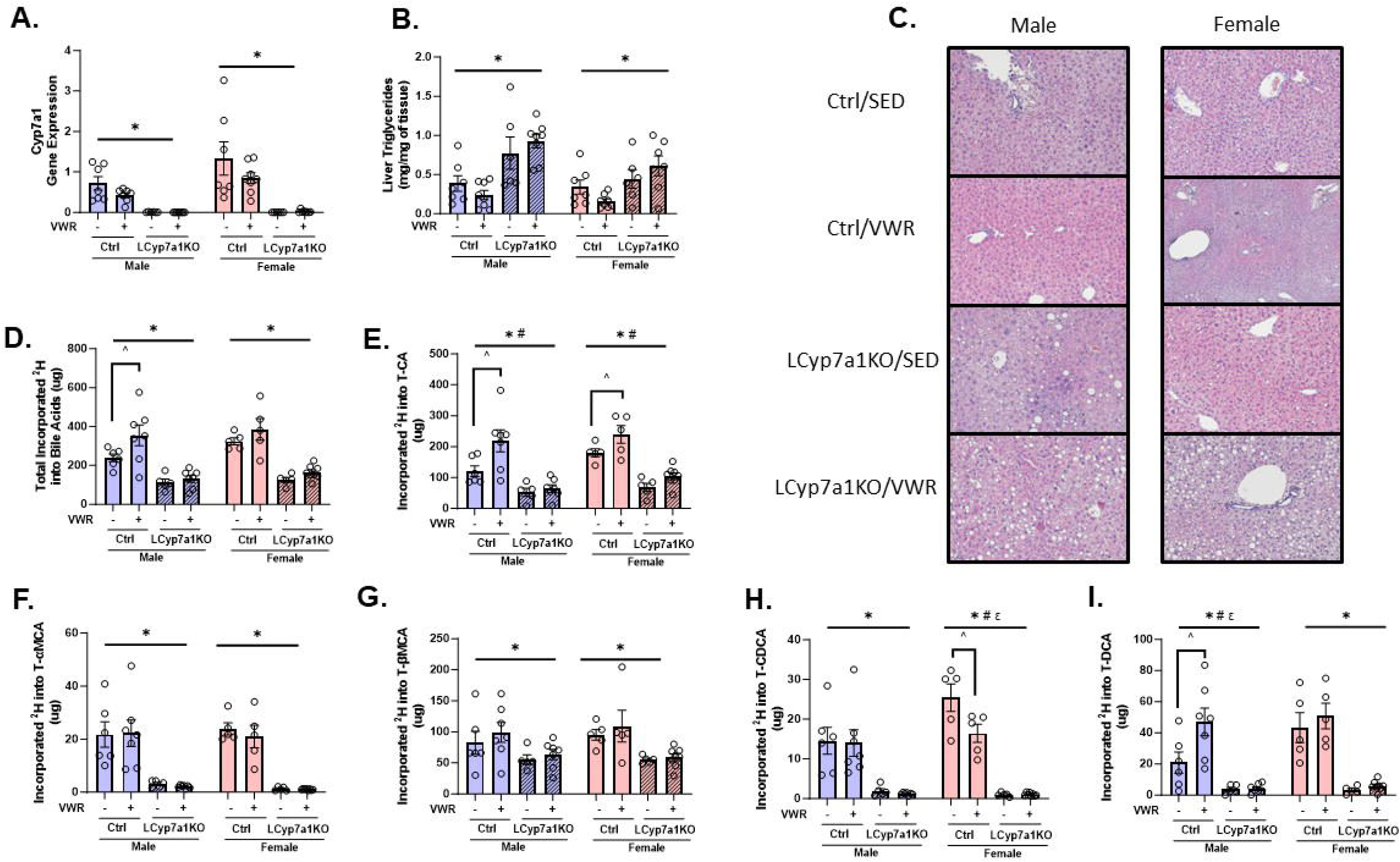
Liver triglyceride and bile acid content in liver-specific Cyp7a1 knockout mice with VWR. **A.** Liver Cyp7a1 gene expression. **B.** Liver triglyceride content. **C.** Representative hematoxylin and eosin stains. **D.** Total liver bile synthesis. **E.** Liver T-CA bile acid synthesis. **F.** Liver T-αMCA bile acid synthesis. **G**. Liver T-βMCA bile acid synthesis. **H.** Liver T-CDCA bile acid synthesis. **I.** Liver T-DCA bile acid synthesis. Data represented as means ± SEM, (n=6-8). * Indicates main effect of LCyp7a1KO within sex (*P<0.05), # indicates main effect of VWR within sex (P<0.05), ε indicates an LCyp7a1KO and VWR interaction within sex, ^ P<0.05 vs. indicated group.

The fraction of new bile acids following ^2^H_2_O administration were not remarkably different, perhaps due to the much smaller pool sizes in the LCyp7a1KO mice, but the absolute amounts of new bile acids in LCyp7a1KO mice were substantially reduced, consistent with impaired bile acid synthesis (Main effect of LCyp7a1KO, P<0.05, **Fig 6D**). This reduction was evident across multiple bile acid species, including T-CA, T-αMCA, T-βMCA, T-CDCA, and T-DCA, regardless of sex (P<0.05, Main effect of LCyp7a1KO, **Fig 6E-I**). Notably, exercise was associated with an upregulation of bile acid synthesis in both male (47.9%) and female (18.9%) control mice, an effect that was absent in LCyp7a1KO mice (P<0.05, **Fig 6D**). This was primarily driven by an increase in T-CA with exercise in male (81.1%) and female (32.8%) control mice (P<0.05, **Fig 6E**). Additionally, male control mice that exercised exhibited increased T-DCA bile acid synthesis (P<0.05, **Fig 6I**). In contrast, female control mice that exercised showed a reduction in T-CDCA bile acid synthesis (P<0.05, **Fig 6H**). These data indicate that bile acid synthesis through *Cyp7a1* plays an important role in exercise-mediated protection from diet-induced liver steatosis.

## 4. DISCUSSION

Higher aerobic capacity and exercise are known to prevent and treat metabolic diseases, including MASLD [5–8], respectively. We previously reported that higher aerobic capacity and exercise enhance hepatic gene expression of the bile acid pathway and increase fecal bile acid loss in rodents [10; 11]. Moreover, a previous study reported that chronic exercise increased fecal bile acid excretion, accompanied by increased bile acid flow and biliary secretion of cholate-derived bile acids [12]. However, whether hepatic bile acid synthesis is elevated by exercise and if this adaptation plays a critical role in liver metabolism, including the treatment of hepatic steatosis, remained unclear. To assess *in vivo* bile acid synthesis, we administered ^2^H_2_O and tracked ^2^H incorporation into bile acids by LC-MS/MS detection. These data confirmed that HCR rats have higher bile acid synthesis than LCR rats. Furthermore, 4 weeks of exercise increased hepatic bile acid synthesis in wild-type mice. Consistent with our previous research, both higher aerobic capacity and exercise upregulated *Cyp7a1* gene expression, suggesting that *Cyp7a1* may be essential for the metabolic benefits of both intrinsic exercise capacity and daily physical exercise. For the first time, we also show that the knockout of hepatic *Cyp7a1* reduced bile acid content but increased hepatic steatosis and that it negated the capacity of exercise to lower hepatic steatosis induced by a chronic HFD. Overall, the data show that the regulation of *Cyp7a1* and bile acid synthesis play a critical role in aerobic capacity and exercise ability in combating MASLD.

Metabolic flexibility, or the capacity to efficiently switch between fuel sources depending on nutrient availability, is crucial for maintaining metabolic health. Impaired metabolic flexibility, such as the inability to properly regulate hepatic lipid synthesis and/or oxidation, is strongly associated with insulin resistance and hepatic steatosis [35–37], while multiple lines of evidence show that exercise improves metabolic flexibility [38]. Our previous studies demonstrated that HCR rats are protected from HFD-induced insulin resistance and hepatic steatosis and provided evidence of pronounced differences in their whole-body metabolic flexibility, indicated by a superior capacity to upregulate dietary FAO when transitioned to a high-fat diet [7; 39]. However, no studies have assessed the capacity of HCR and LCR rat models to moderate DNL in response to nutritional conditions. Consistent with previous research in mice and rats [8; 33], we observed that DNL was stimulated in the fed state and was highest on the carbohydrate-rich LFD. Interestingly, HCR rats displayed a more robust induction of DNL on a LFD, and they suppressed DNL more completely on a HFD compared to LCR rats. The heightened metabolic flexibility of DNL in HCR livers may contribute to their exceptional metabolic profile, such as improved glycemia during high carbohydrate consumption, by increasing the disposal of glucose carbons into lipid stores, or reduced hepatic steatosis during high fat consumption, by activating fat oxidation with obligate inhibition of DNL. Likewise, similar factors may also play a role in the upregulation of cholesterol and bile acid synthesis in HCR when fed a HFD for 1 week. The shunting of cytosolic acetyl-CoA towards cholesterol and bile acid synthesis may contribute to lower DNL in HCR rats on a HFD. Since sterol synthesis does not require malonyl-CoA, a potent inhibitor of mitochondrial fat transport and oxidation, its increased activity may preserve FAO. Indeed, FAO and mitochondrial respiration are increased in HCR rats [10; 26], which may also facilitate the energy-costly cholesterol and bile acid synthesis pathways. Mechanistic studies will need to be undertaken to test the precise link between the activation of bile acid synthesis and increased metabolic flexibility endowed by exercise or intrinsic aerobic capacity.

Our findings reveal a novel link between aerobic capacity, exercise, cholesterol, and bile acid synthesis. Our data shows that HCR rats have enhanced cholesterol synthesis despite maintaining lower serum cholesterol levels, particularly after prolonged HFD feeding. This observation suggests an increased channeling of cholesterol towards bile acid synthesis and fecal excretion in HCR. HCR rats consistently display greater fecal bile acid loss, aligning with previous research in exercising mice demonstrating elevated bile acid excretion and cholesterol turnover that was previously linked to increased survival and reduced atherosclerotic lesions in LDL-R knockout mice [12; 40]. Chronic exercise in mice also upregulates fecal bile acid loss and tracer studies demonstrate a concomitant increase in bile acid synthesis. These findings are further supported by our previous work in both rodents and humans, where we observed a consistent pattern of increased fecal bile acid levels and/or enhanced expression of hepatic genes involved in cholesterol and bile acid metabolism in response to exercise training [11; 41]. Moreover, we showed that improving fitness and reducing body weight with a diet and exercise intervention in middle-aged, obese women increased a known marker of bile acid synthesis (C4), while also appearing to enhance bile acid feedback regulation [42]. In a previous study, we also compared markers of bile acid metabolism in women with high aerobic capacity vs. moderate aerobic capacity matched for body weight and age [41]. That study did not reveal differences in markers of bile acid synthesis or fecal excretion, likely due to dietary controls that induced unintentional weight loss in high-fit women with very high daily activity levels. However, notably, a marker of bile acid synthesis (C4) and bile acid species were markedly different between high and moderate-fit women during postprandial conditions (OGTT). Glucose and insulin are known regulators of *Cyp7a1* expression and bile acid metabolism, suggesting that aerobic capacity not only regulates bulk bile acid synthesis but also may modulate a sophisticated regulation of bile acid metabolism right after feeding.

Collectively, our data in rodents suggest that higher aerobic capacity and exercise promote a shift in cholesterol metabolism toward increased bile acid synthesis and fecal excretion, which appear to facilitate some beneficial effects of exercise on liver health. The primary mechanisms of action by which fitness or exercise leads to greater *Cyp7a1*-mediated bile acid synthesis are unknown but could be linked to higher intestinal motility or less bile acid absorption in the intestines or colon, leading to greater fecal bile acid loss and commensurate increases in bile acid synthesis to maintain homeostasis. However, exercise-induced changes in bile acid metabolism may also result from primary changes in production. A previous study using a crossover within-subject design reported that bile acid levels in the duodenum increased by 10-fold following 30 min of light-intensity exercise vs. sedentary conditions in young men, despite no large difference in total fluid in the duodenum or changes in gall bladder size [43]. This finding could suggest that each bout of exercise increases the production of bile acids, and thus, turnover increases with fecal excretion rising as a result. A newer study found that acute resistance exercise and acute endurance exercise both lowered circulating bile acid levels and increased a bile acid species known to target TGR5 receptors, lithocholic acid, but differently regulated circulating levels of FGF19 and FGF21, which play a role in feedback regulation of bile acid synthesis in the liver [44]. However, the effects of acute exercise on bile acid metabolism do not explain the divergent HCR vs. LCR phenotype occurring in rats maintained in a sedentary condition. Exercise and aerobic capacity also sensitize hepatic insulin signaling, which potently upregulates *CYP7A1* enzyme expression. Thus, differences in the capacity of insulin to upregulate *Cyp7a1* and bile acid synthesis, in addition to regulating shuttling of acetyl CoA away from DNL towards bile acid synthesis may also play a causal role between high and low aerobic capacity or exercise vs. sedentary conditions. Differences in insulin sensitivity can also influence bile acid pool composition through the enzyme *CYP8B1* [45]. *CYP8B1* is an enzyme in the bile acid synthetic pathway responsible for the 12-alpha hydroxylation of bile acids and, therefore, determines the 12-alpha to non-12-alpha hydroxylated bile acid ratio. Insulin action suppresses *CYP8B1* activity through the nuclear exclusion of *FOXO1*, however, insulin resistance causes the ratio of 12-alpha to non-12alpha hydroxylated bile acids to increase [46]. After 20 weeks of a HFD, this ratio was much higher in LCR rats than HCR rats, consistent with our previous observation of worsening metabolic health and reduced insulin signaling in LCR rats on a chronic HFD [6]. There was no significant difference in 12-alpha to non-12-alpha hydroxylated bile acids in the 1-week study, suggesting that initial insulin signaling differences between the strains are not a factor. In contrast, alterations in *Cyp27a1* and *Cyp7b1*, suggest an upregulation of the non-12-alpha hydroxylated bile acid, CDCA, pathway. *Cyp27a1* and *Cyp7b1* are the main regulatory steps in this alternative bile acid synthetic pathway [47]. *Cyp27a1* is localized in mitochondria and is responsible for the side-chain oxidation needed to form bile acids in both the classic and alternative pathways [48]. Hence, the increased expression of *Cyp27a1* in HCR liver likely contributes to a higher overall bile acid synthesis rate and is consistent with our previous finding of higher hepatic *Pgc1α* expression and mitochondrial content in HCR liver [49]. In contrast, greater expression of *Cyp7b1* is a more specific indication that the alternative bile acid pathway is upregulated in LCR liver.

Bile acid synthesis occurs via two pathways: classic and alternative. Overexpression of *Cyp7a1*, a rate-limiting enzyme in the classic pathway, attenuates weight gain on a HFD and improves metabolic health, including protecting against hepatic steatosis [50]. Consistent with this, HCR rats and exercise upregulate *Cyp7a1* expression, suggesting a potential role for the classic pathway in preventing and treating hepatic steatosis. However, the relationship between *Cyp7a1* and hepatic steatosis is complex. While *Cyp7a1*-deficient mice from birth exhibit protection from metabolic disorders without altering hepatic steatosis on a HFD [51]. However, bile acids are critical for the digestion and absorption of lipids, and the *Cyp7a1* knockout model reportedly displayed a leanness phenotype due to an inability to digest dietary lipids. In contrast, in this study, the inducible liver-specific *Cyp7a1* knockout model displayed normal weight on the HFD compared to controls and developed increased hepatic steatosis in both sexes. This discrepancy between the knockout methodologies may arise from the reduced capacity of the alternative pathway of *Cyp27a1* to compensate for *Cyp7a1* deficiency in our model or from the fact that we allowed *Cyp7a1* to be functional past a critical developmental window.

## 5. CONCLUSION

In conclusion, this study provides novel insights into the link between aerobic capacity, exercise, bile acid metabolism, and steatosis. Our findings demonstrate that both intrinsic high aerobic capacity and exercise training enhance bile acid synthesis. Elevated bile acid synthesis, driven by *Cyp7a1*, appears critical for the beneficial effects of exercise to treat steatosis induced by a HFD. Importantly, our results identify bile acid synthesis as a key mediator between aerobic capacity, exercise, and hepatic energy metabolism that may also be linked to whole-body metabolism and long-term risk for type 2 diabetes and MASLD, which have shown to be independently linked to aerobic capacity and exercise behavior in human studies. Further investigation is warranted to understand the mechanisms of action by which intrinsic aerobic capacity and exercise lead to greater bile acid synthesis.

## Abbreviations

(VWR): Voluntary wheel running
(BA): bile acids
(HCR): high capacity runner rats
(LCR): low capacity runner rats
(LFD): low fat diet
(HFD): high fat diet
(MASLD): Metabolic dysfunction-associated steatotic liver disease
(FAO): Fatty acid oxidation

## DATA AVAILABILITY STATEMENT

All data are found within the manuscript.

## ACKNOWLEDGMENTS

We thank Drs. Greg Graf, Udayan Apte, and E. Matthew Morris, for their intellectual contributions to previous findings that proceeded with this work. We thank Samantha J. McKee at the University of Toledo for expert phenotyping, care, and maintenance of the LCR/HCR rat colony.

## CRediT

**BAK**: Writing – Original Draft, Conceptualization, Formal Analysis, Investigation, Data Curation, Project Administration. **AM**: Writing – Original Draft, Conceptualization, Formal Analysis, Investigation, Data Curation, Project Administration. **XF**: Writing-Review and editing, Methodology, Formal Analysis, Investigation, Data Curation. **EF**: Data Curation, Investigation, **NE**: Data Curation. **KS**: Data Curation. **JA**: Data Curation. **TL**: Conceptualization, Methodology. **LK**: Conceptualization, Methodology. **SB** Conceptualization, Methodology. **PC:** Methodology, Writing-Review and editing; **SB:** Conceptualization, Funding Acquisition, Methodology, Supervision, Writing-Review and editing. **JT:** Conceptualization, Funding Acquisition, Methodology, Supervision, Writing-Review and editing.

---

**Supplemental Figure 1.** Representative LC-MS/MS chromatogram of rat liver tissue. Structural isomers, T-αMCA, T-βMCA, and T-CA were completely separated. T-UDCA, T-CDCA, and T-DCA isomers were baseline-separated as well.

**Supplemental Figure 2.** Bile acid content in liver-specific Cyp7a1 knockout mice with VWR. **A.** Liver bile content. **B.** Gallbladder bile acid content. **C.** Intestine bile acid content. **D**. Fecal bile acid content. Data represented as means ± SEM, (n=6-8). *indicates main effect of LCyp7a1 within sex (*P<0.05).

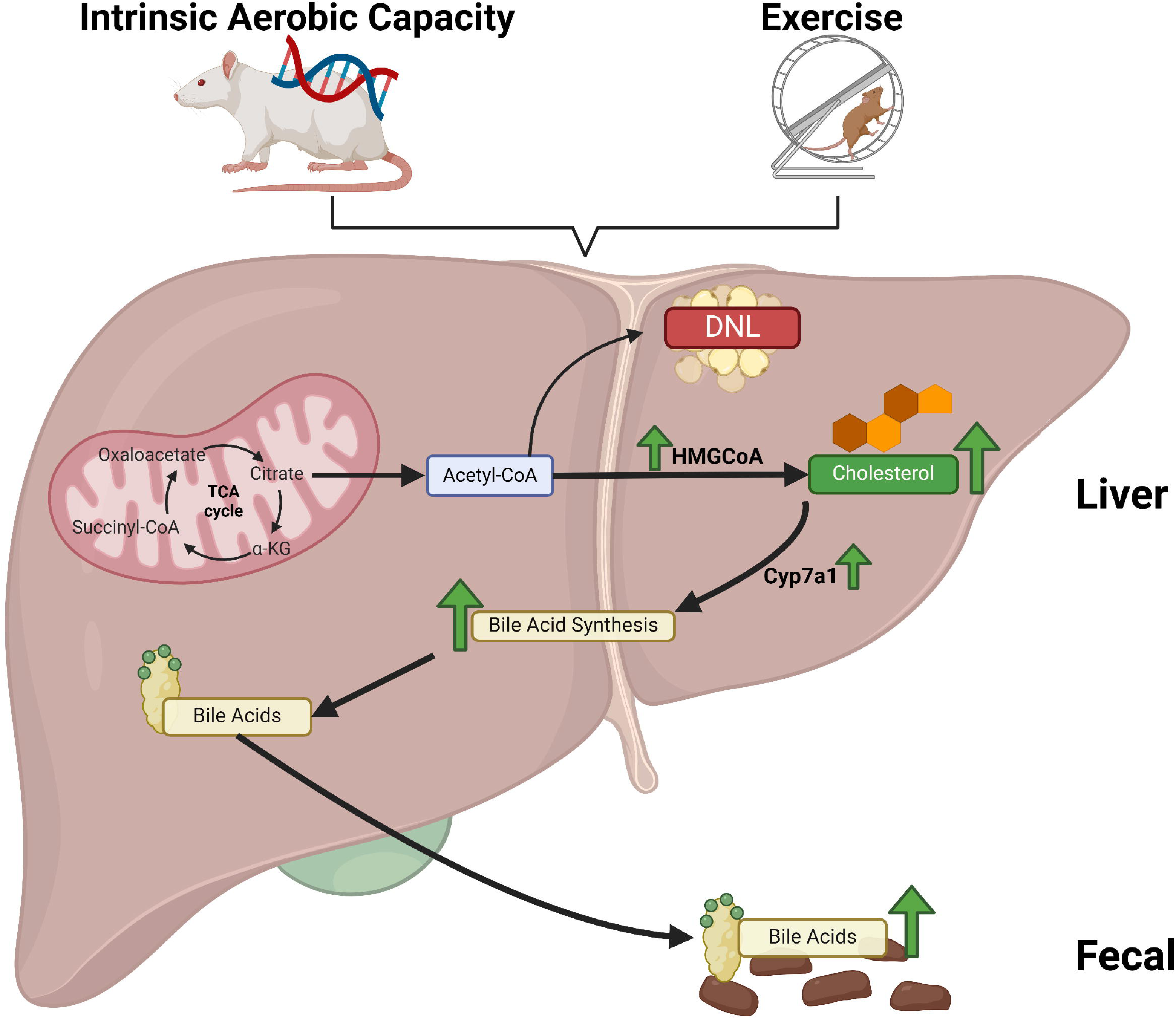

