## Supplemental Tables for "Aerobic capacity and exercise mediate protection against hepatic steatosis via enhanced bile acid metabolism"

**Supplemental Table S1.** MRM transition of Bas and deuterium-labeled internal standards

| **Compound** | **MID** | **Q1 m/z** | **Q3 m/z** |
|---|---|---|---|
| TMCA, TCA | M+0 | 514.2 | 80.0 |
| | M+1 | 515.2 | 80.0 |
| | M+2 | 516.2 | 80.0 |
| | M+3 | 517.2 | 80.0 |
| TCDCA, TDCA | M+0 | 498.2 | 80.0 |
| | M+1 | 499.2 | 80.0 |
| | M+2 | 500.2 | 80.0 |
| | M+3 | 501.2 | 80.0 |
| d9-TCDCA | | 507.2 | 80.0 |

**Supplemental Table S2.** Anthropometric and energy intake data from HCR/LCR rats on a LFD or HFD for 1-week.

| | LCR | | HCR | | P-value | | |
|---|---|---|---|---|---|---|---|
| **Variable** | **LFD** | **HFD** | **LFD** | **HFD** | **Strain** | **Diet** | **Strain x Diet** |
| ∆ Body Mass (g) | 8.9 ± 1.7 | 26.6 ± 2.2*** | 7.1 ± 2.3 | 15.8 ± 1.4*,^ | **0.0003** | **0.001** | **0.027** |
| ∆ Fat Mass (g) | 2.3 ± 1.0 | 19 ± 2.3*** | 6.2 ± 1.5 | 9.7 ± 2.2 | 0.155 | **0.001** | **0.001** |
| ∆ Lean Mass (g) | -1.3 ± 1.2 | -2.5 ± 2.4 | -7.5 ± 1.6 | -1.6 ± 1.5 | 0.135 | 0.187 | 0.052 |
| Body Mass (g) | 452 ± 14.8 | 480 ± 15.4 | 350 ± 14.1^^^ | 356 ± 8.9^^^ | **<0.001** | 0.213 | 0.445 |
| % Fat Mass | 15.9 ± 1.0 | 19.7 ± 1.1* | 9.7 ± 0.5^^^ | 11.6 ± 1.0^^^ | **<0.001** | **0.006** | 0.342 |
| % Lean Mass | 76.6 ± 0.9 | 73 ± 1.1* | 83.1 ± 0.5^^^ | 81.1 ± 1.1^^^ | **<0.001** | **0.005** | 0.355 |
| Total Energy Intake (kcal) | 418 ± 18.6 | 612 ± 35.1*** | 460 ± 28.6 | 524 ± 14.4 | 0.377 | **0.001** | **0.016** |

Values are means ± SEM (n=8; Only rats maintaining access to food prior to sacrifice). Significance was determined by 2-way ANOVA (strain X diet) followed by Tukey's multiple comparisons test; *indicates effect of diet within strain (*p<0.05, **p<0.01, ***p<0.001); ^indicates effect of strain within diet (^p<0.05, ^^p<0.01, ^^^p<0.001).

**Supplemental Table S3.** Anthropometric and energy intake data from HCR/LCR rats on a LFD or HFD for 20-weeks.

| | LCR | | HCR | | P-value | | |
|---|---|---|---|---|---|---|---|
| **Variable** | **LFD** | **HFD** | **LFD** | **HFD** | **Strain** | **Diet** | **Strain x Diet** |
| Δ Body Mass (g) | 75.3 ± 12.9 | 107 ± 12.7 | 69.3 ± 6.6 | 89.9 ± 8.1 | 0.27 | **0.016** | 0.586 |
| Body Mass (g) | 506 ± 16.5 | 541.0 ± 13.0 | 411 ± 11.9^^^ | 426.0 ± 20.7^^^ | **<0.001** | 0.117 | 0.533 |
| % Fat Mass | 19.8 ± 2.3 | 25.9 ± 1.7 | 16.0 ± 1.0 | 18.6 ± 1.4^ | **0.002** | **0.013** | 0.298 |
| % Lean Mass | 72.7 ± 2.1 | 67.0 ± 1.7 | 76.1 ± 1.1 | 73.9 ± 1.3^ | **0.003** | **0.018** | 0.285 |
| Weekly Energy Intake (kcal) | 406 ± 8.3 | 429 ± 7.1 | 389 ± 10.2 | 407 ± 13.8 | 0.058 | 0.052 | 0.833 |

Values are means ± SEM (n=10). Significance was determined by 2-way ANOVA (strain X diet) followed by Tukey's multiple comparisons test; †indicates diet effect (†p<0.05, ††p<0.01, †††p<0.001); ^indicates effect of strain within diet (^p<0.05, ^^p<0.01, ^^^p<0.001).

**Supplemental Table S4.** Serum metabolic data from HCR/LCR rats on a LFD or HFD for 1-week.

| | **LCR** | | | | **HCR** | | | | **P-value** | | | | |
|---|---|---|---|---|---|---|---|---|---|---|---|---|---|
| | **FASTED** | | **FED** | | **FASTED** | | **FED** | | | | | | |
| **Variable** | **LFD** | **HFD** | **LFD** | **HFD** | **LFD** | **HFD** | **LFD** | **HFD** | **Strain** | **FED** | **Diet** | **Strain x FED** | **Strain x Diet** |
| ALP (U/L) | 64.5 ± 5.6 | 88.1 ± 8.6 | 131 ± 9.1‡‡‡ | 235 ± 15.8‡‡‡,*** | 106 ± 6.4 | 105 ± 7.5 | 179 ± 5.2^,‡‡‡ | 253 ± 18.0‡‡‡,*** | **<0.001** | **<0.001** | **<0.001** | 0.825 | 0.071 |
| AST (U/L) | 118 ± 12.4 | 124 ± 4.7 | 149.0 ± 7.0 | 142 ± 8.3 | 128 ± 6.3 | 132 ± 16.7 | 277 ± 74.8‡ | 175.0 ± 22.1 | **0.034** | **0.005** | 0.238 | 0.087 | 0.239 |
| ALT(U/L) | 55.6 ± 9.7 | 52.8 ± 4.6 | 62.0 ± 6.4 | 65.1 ± 3.5 | 69.9 ± 3.9 | 69.3 ± 6.3 | 183 ± 69.3 | 116 ± 32.1 | **0.012** | **0.025** | 0.393 | 0.074 | 0.389 |
| Albumin (g/dL) | 2.8 ± 0.0 | 2.9 ± 0.1 | 3.3 ± 0.0‡‡‡ | 3.2 ± 0.0‡‡‡ | 2.8 ± 0.0 | 2.7 ± 0.0 | 3.3 ± 0.1‡‡‡ | 3.3 ± 0.0‡‡‡ | 0.801 | **<0.001** | 0.675 | 0.082 | 0.451 |
| Total Protein (g/dL) | 4.9 ± 0.1 | 5.2 ± 0.1 | 5.9 ± 0.1‡‡‡ | 5.9 ± 0.1‡‡‡ | 4.9 ± 0.1 | 4.8 ± 0.1 | 6.0 ± 0.2‡‡‡ | 5.9 ± 0.1‡‡‡ | 0.244 | **<0.001** | 0.716 | 0.161 | 0.102 |
| BUN (mg/dL) | 11.5 ± 0.8 | 10.8 ± 0.9 | 14.1 ± 0.4 | 16.3 ± 0.9‡‡‡ | 11.6 ± 0.8 | 9.1 ± 0.3 | 17.3 ± 0.7‡‡‡ | 13.8 ± 0.5‡‡‡,* | 0.657 | **<0.001** | **0.022** | 0.284 | **<0.001** |
| Cholesterol (mg/dL) | 68.0 ± 6.2 | 10.8 ± 7.7 | 94.4 ± 3.3‡ | 16.3 ± 6.6‡ | 57.1 ± 5.0 | 9.1 ± 5.3 | 75.8 ± 5.2 | 13.8 ± 5.4^ | **<0.001** | **<0.001** | **0.042** | 0.337 | 0.096 |
| Glucose (mg/dL) | 159 ± 6.5 | 156 ± 11.8 | 259 ± 16.5‡‡‡ | 250 ± 7.1‡‡‡ | 154 ± 8.4 | 162 ± 6.8 | 258 ± 20.1‡‡‡ | 269 ± 17.0‡‡‡ | 0.591 | **<0.001** | 0.899 | 0.648 | 0.394 |
| Triglycerides (mg/dL) | 94.8 ± 11.1 | 104 ± 25.0 | 93 ± 5.1 | 142 ± 19.7 | 57.9 ± 6.8 | 50.0 ± 3.7 | 112 ± 19.1 | 92.4 ± 14.1^ | **0.006** | **0.003** | 0.465 | 0.160 | **0.049** |
| β-Hydroxybutyrate (mg/dL) | 11.4 ± 0.8 | 14.4 ± 1.7 | 2.6 ± 0.2‡‡‡ | 3.4 ± 0.2‡‡‡ | 11.6 ± 1.1 | 15.8 ± 2.0 | 2.4 ± 0.2‡‡‡ | 3.1 ± 0.3‡‡‡ | 0.718 | **<0.001** | **0.005** | 0.482 | 0.711 |
| NEFA (mEq/L) | 0.13 ± 0.01 | 0.15 ± 0.02 | 0.2 ± 0.02 | 0.22 ± 0.03 | 0.15 ± 0.02 | 0.14 ± 0.02 | 0.11 ± 0.01 | 0.12 ± 0.03 | **0.003** | 0.143 | 0.548 | **0.001** | 0.391 |
| Insulin (ng/mL) | 3.2 ± 0.3 | 4.2 ± 0.7 | 3.9 ± 1.3 | 4.7 ± 1.5 | 2.9 ± 1.1 | 5.3 ± 1.0 | 7.2 ± 1.8 | 9.0 ± 2.6 | **0.042** | **0.031** | 0.152 | 0.099 | 0.575 |

Values are means ± SEM. Significance was determined by 3-way ANOVA (strain X diet X fed state) followed by Tukey's multiple comparisons test; *indicates effect of diet within strain and fed state (*p<0.05, **p<0.01, ***p<0.001); ^indicates effect of strain within diet and fed state (^p<0.05, ^^p<0.01, ^^^p<0.001); ‡indicates effect of fed state within strain and diet (‡p<0.05, ‡‡p<0.01, ‡‡‡p<0.001).

**Supplemental Table S5.** Serum metabolic data from HCR/LCR rats on a LFD or HFD for 20-weeks.

| | LCR | | | | HCR | | | | P-value | | |
|---|---|---|---|---|---|---|---|---|---|---|---|
| **Variable** | **LFD** | | **HFD** | | **LFD** | | **HFD** | | **Strain** | **Diet** | **Strain x Diet** |
| ALP (U/L) | 161.1 | ± 12.9 | 230.9 | ± 15.6* | 189.9 | ± 13.1 | 231.0 | ± 18.5 | 0.348 | **<0.001** | 0.351 |
| AST (U/L) | 156.7 | ± 16.3 | 180.9 | ± 13.7 | 169.7 | ± 13.6 | 148.1 | ± 19.4 | 0.538 | 0.935 | 0.159 |
| ALT (U/L) | 87.2 | ± 11.0 | 113.0 | ± 11.9 | 124.9 | ± 18.9 | 107.7 | ± 22.6 | 0.342 | 0.800 | 0.210 |
| Albumin (g/dL) | 3.3 | ± 0.1 | 3.3 | ± 0.0 | 3.3 | ± 0.1 | 3.1 | ± 0.1 | 0.305 | 0.137 | 0.238 |
| Total Protein (g/dL) | 6.0 | ± 0.1 | 6.0 | ± 0.1 | 6.1 | ± 0.1 | 5.7 | ± 0.1 | 0.418 | **0.044** | 0.110 |
| BUN (mg/dL) | 16.5 | ± 0.6 | 15.3 | ± 0.6 | 16.0 | ± 0.7 | 14.3 | ± 1.1 | 0.342 | 0.071 | 0.750 |
| Cholesterol (mg/dL) | 122.6 | ± 4.8 | 116 | ± 4.9 | 97.0 | ± 4.5^^ | 93.3 | ± 3.5^^ | **<0.001** | 0.255 | 0.747 |
| Glucose (mg/dL) | 220.4 | ± 8.5 | 203.3 | ± 6 | 247.4 | ± 14.7 | 213.9 | ± 7.9 | 0.065 | **0.015** | 0.411 |
| Triglycerides (mg/dL) | 77.3 | ± 5.9 | 134.5 | ± 7.2** | 137.6 | ± 12.0^^ | 109.2 | ± 17.2 | 0.135 | 0.216 | **<0.001** |
| β-Hydroxybutyrate (mg/dL) | 2.6 | ± 0.2 | 3.7 | ± 0.3 | 2.3 | ± 0.1 | 4.0 | ± 0.3 | 0.913 | **<0.001** | 0.199 |
| NEFA (mEq/L) | 0.26 | ± 0.04 | 0.55 | ± 0.04** | 0.4 | ± 0.07 | 0.49 | ± 0.07 | 0.474 | **0.002** | 0.083 |

Values are means ± SEM. Significance was determined by 2-way ANOVA (strain X diet) followed by Tukey's multiple comparisons test; *indicates effect of diet within strain (*p<0.05, **p<0.01, ***p<0.001); ^indicates effect of strain within diet (^p<0.05, ^^p<0.01, ^^^p<0.001).

**Supplemental Table S6.** Serum bile acid data from HCR/LCR rats on a LFD or HFD for 1-week.

| (ng/mL) | LCR | | | | HCR | | | | P-value | | | | |
|---|---|---|---|---|---|---|---|---|---|---|---|---|---|
| | FASTED | | FED | | FASTED | | FED | | | | | | |
| | LFD | HFD | LFD | HFD | LFD | HFD | LFD | HFD | Strain | FED | Diet | Strain x FED | Strain x Diet |
| G-CA | 229.41 ± 57.24 | 86.15 ± 30.81** | 10.93 ± 4.20‡ | 16.14 ± 6.01 | 32.26 ± 12.00^^ | 11.92 ± 3.13 | 4.05 ± 1.43 | 5.24 ± 1.43 | **<0.001** | **<0.001** | **0.022** | **<0.001** | 0.080 |
| G-CDCA | 12.08 ± 4.49 | 4.41 ± 1.90 | 0.34 ± 0.10‡‡‡ | 0.82 ± 0.38 | 1.32 ± 0.55^^ | 0.52 ± 0.19 | 0.33 ± 0.12 | 0.24 ± 0.11 | **0.003** | **0.001** | 0.105 | **0.006** | 0.210 |
| G-DCA | 29.41 ± 15.51 | 7.83 ± 4.04 | 0.08 ± 0.06‡ | 0.12 ± 0.14 | 3.23 ± 1.31^ | 1.05 ± 0.19 | 0.08 ± 0.09 | -0.01 ± 0.06 | **0.045** | **0.014** | 0.146 | **0.045** | 0.237 |
| G-UDCA | 0.58 ± 0.10 | 0.14 ± 0.08*** | 0.04 ± 0.05‡‡‡ | 0.13 ± 0.10 | 0.03 ± 0.05^^^ | 0.04 ± 0.04 | -0.05 ± 0.03 | -0.02 ± 0.03 | **<0.001** | **0.003** | **0.024** | **0.022** | 0.059 |
| T-αMCA | 69.66 ± 8.82 | 49.35 ± 6.98 | 83.23 ± 26.17 | 144.86 ± 45.36 | 53.46 ± 6.99 | 48.69 ± 12.89 | 28.15 ± 8.26 | 38.19 ± 10.81^^ | **0.003** | 0.205 | 0.419 | **0.014** | 0.531 |
| T-βMCA | 24.04 ± 8.84 | 29.82 ± 9.14 | 174.74 ± 67.21 | 200.30 ± 81.98 | 48.70 ± 13.80 | 77.76 ± 17.14 | 27.04 ± 8.62 | 75.18 ± 29.11 | 0.082 | **0.011** | 0.342 | **0.004** | 0.687 |
| T-CA | 344.11 ± 88.20 | 226.07 ± 55.23 | 411.42 ± 155.45 | 772.35 ± 277.73 | 250.66 ± 58.79 | 350.10 ± 78.81 | 120.35 ± 50.78 | 301.04 ± 85.62 | **0.049** | 0.237 | 0.155 | **0.033** | 0.919 |
| T-CDCA | 18.01 ± 2.72 | 11.74 ± 1.80 | 17.45 ± 4.83 | 57.71 ± 22.19‡ | 14.21 ± 2.80 | 15.66 ± 4.47 | 15.51 ± 5.74 | 18.61 ± 4.94 | 0.104 | **0.049** | 0.125 | 0.102 | 0.239 |
| T-DCA | 47.84 ± 27.19 | 19.29 ± 4.72 | 12.12 ± 3.00 | 19.32 ± 8.79 | 36.41 ± 12.75 | 29.95 ± 4.07 | 6.60 ± 2.47 | 8.26 ± 2.35 | 0.592 | **0.009** | 0.421 | 0.626 | 0.610 |
| T-LCA | 1.37 ± 0.14 | 0.88 ± 0.19 | 0.28 ± 0.14 | 0.15 ± 0.19 | 1.40 ± 0.91 | 0.75 ± 0.29 | 0.14 ± 0.19 | 0.16 ± 0.11 | 0.833 | **0.001** | 0.207 | 0.759 | 0.617 |
| T-UDCA | 2.34 ± 0.75 | 1.96 ± 0.53 | 10.68 ± 4.02 | 27.67 ± 12.45‡‡ | 3.16 ± 9.02 | 4.17 ± 1.05 | 3.41 ± 1.51 | 4.37 ± 1.27^ | **0.044** | **0.013** | 0.170 | 0.015 | 0.278 |
| αMCA | 38.24 ± 9.83 | 33.01 ± 15.35 | 18.17 ± 11.59 | 23.54 ± 15.06 | 22.61 ± | 10.43 ± 5.08 | 40.50 ± 21.43 | 9.05 ± 2.93 | 0.398 | 0.715 | 0.228 | 0.201 | 0.224 |
| βMCA | 43.16 ± 8.72 | 39.93 ± 15.38 | 73.82 ± 44.36 | 50.44 ± 26.66 | 64.81 ± 16.89 | 32.68 ± 6.54 | 60.05 ± 28.80 | 16.69 ± 5.55 | 0.611 | 0.753 | 0.120 | 0.342 | 0.453 |
| CA | 436.24 ± 118.84 | 208.96 ± 81.73 | 139.03 ± 96.60 | 105.48 ± 64.59 | 104.21 ± 38.54^ | 2.61 ± 0.86 | 96.01 ± 44.12 | 25.52 ± 14.65 | **0.001** | 0.053 | **0.030** | **0.037** | 0.651 |
| CDCA | 63.02 ± 19.72 | 60.12 ± 34.28 | 10.05 ± 7.53 | 17.96 ± 11.88 | 22.63 ± 10.24 | 0.99 ± 0.86 | 28.18 ± 12.18 | 5.48 ± 4.83 | **0.045** | 0.065 | 0.391 | **0.023** | 0.278 |
| DCA | 41.62 ± 21.00 | 17.85 ± 3.85 | 9.44 ± 4.09 | 6.95 ± 1.81 | 33.55 ± 7.33 | 31.60 ± 6.85 | 8.30 ± 2.61 | 8.89 ± 1.87 | 0.790 | **<0.001** | 0.259 | 0.841 | 0.309 |
| LCA | 5.71 ± 0.63 | 5.41 ± 0.90 | 6.08 ± 1.23 | 2.97 ± 0.61 | 7.92 ± 1.79 | 5.34 ± 1.01 | 4.07 ± 0.75 | 2.99 ± 1.07 | 0.960 | **0.008** | **0.022** | 0.174 | 0.933 |
| UDCA | 7.15 ± 1.41 | 7.12 ± 3.20 | 5.65 ± 3.71 | 4.90 ± 2.82 | 7.23 ± 2.03 | 2.40 ± 0.72 | 5.42 ± 2.43 | 1.38 ± 0.77 | 0.220 | 0.338 | 0.155 | 0.884 | 0.232 |
| G-TOTALS | 271.74 ± 71.79 | 98.69 ± 36.27** | 11.31 ± 4.32‡‡‡ | 17.19 ± 6.63 | 36.80 ± 13.82^^ | 13.43 ± 3.45 | 4.33 ± 1.59 | 5.49 ± 1.60 | **<0.001** | **<0.001** | **0.025** | **<0.001** | 0.083 |
| T-TOTALS | 507.38 ± 108.60 | 339.10 ± 74.57 | 709.92 ± 257.71 | 1222.36 ± 440.38 | 408.00 ± 83.39 | 527.07 ± 110.43 | 201.20 ± 71.37 | 445.81 ± 130.16 | **0.039** | 0.163 | 0.215 | **0.018** | 0.973 |
| Uconj TOTALS | 635.13 ± 155.22 | 372.39 ± 137.04 | 262.23 ± 168.31 | 212.23 ± 122.55 | 262.96 ± 78.88 | 86.05 ± 15.53 | 242.54 ± 105.93 | 70.00 ± 29.13 | **0.014** | 0.084 | **0.046** | 0.131 | 0.910 |
| % G-TOTALS | 19.43 ± 3.92 | 11.03 ± 3.69 | 1.52 ± 0.64‡‡‡ | 1.22 ± 0.09‡ | 4.37 ± 1.10^^^ | 2.15 ± 0.40^ | 0.85 ± 0.22 | 1.02 ± 0.24 | **<0.001** | **<0.001** | 0.058 | **<0.001** | 0.237 |
| % T-TOTALS | 38.42 ± 5.67 | 49.19 ± 12.52 | 72.52 ± 9.58 | 83.42 ± 4.05 | 60.40 ± 8.68 | 81.91 ± 3.28 | 58.14 ± 12.56 | 82.79 ± 8.04 | 0.112 | **0.009** | **0.008** | **0.006** | 0.324 |
| % Unconj TOTALS | 42.14 ± 6.30 | 39.78 ± 12.50 | 25.96 ± 9.68 | 15.35 ± 4.05 | 35.23 ± 8.24 | 15.93 ± 3.45 | 41.01 ± 12.55 | 16.19 ± 7.95 | 0.548 | 0.166 | **0.024** | 0.063 | 0.211 |
| 12-hydroxy/non-hydroxy | 4.60 ± 1.70 | 2.89 ± 0.55 | 1.32 ± 0.12‡ | 1.73 ± 0.06 | 1.85 ± 0.28 | 2.14 ± 0.30 | 1.12 ± 0.20 | 2.03 ± 0.25 | 0.075 | **0.006** | 0.956 | 0.060 | 0.185 |
| Total (ng/mL) | 1414.3 ± 278.20 | 810.19 ± 98.93 | 983.46 ± 302.54 | 1451.79 ± 529.76 | 707.77 ± 126.04 | 626.55 ± 118.05 | 448.07 ± 151.12 | 521.30 ± 130.43 | **0.002** | 0.833 | 0.844 | 0.432 | 0.861 |

Values are means ± SEM. Significance was determined by 3-way ANOVA (strain X diet X fed state) followed by Tukey's multiple comparisons test; *indicates effect of diet within strain and fed state (*$p<0.05$, **$p<0.01$, ***$p<0.001$); ^indicates effect of strain within diet and fed state (^$p<0.05$, ^^$p<0.01$, ^^^$p<0.001$); ‡indicates effect of fed state within strain and diet (‡$p<0.05$, ‡‡$p<0.01$, ‡‡‡$p<0.001$).

**Supplemental Table S7.** Serum bile acid data from HCR/LCR rats on a LFD or HFD for 20-weeks.

| (ng/mL) | LCR | | HCR | | P-value | | |
|---|---|---|---|---|---|---|---|
| | **LFD** | **HFD** | **LFD** | **HFD** | **Strain** | **Diet** | **Strain x Diet** |
| G-CA | 25.00 ± 7.83 | 22.28 ± 4.73 | 11.16 ± 3.67 | 11.38 ± 4.34 | **0.028** | 0.818 | 0.787 |
| G-CDCA | 0.65 ± 0.16 | 0.42 ± 0.09 | 0.37 ± 0.10 | 0.14 ± 0.07 | **0.023** | 0.062 | 0.965 |
| G-DCA | 0.59 ± 0.39 | 0.38 ± 0.06 | 0.07 ± 0.04 | 0.12 ± 0.05 | 0.056 | 0.688 | 0.581 |
| G-UDCA | 0.14 ± 0.05 | 0.05 ± 0.03 | 0.05 ± 0.04 | 0.03 ± 0.03 | 0.152 | 0.231 | 0.629 |
| T-αMCA | 62.16 ± 9.57 | 79.38 ± 22.60 | 48.16 ± 9.96 | 37.05 ± 11.86 | 0.060 | 0.834 | 0.336 |
| βMCA | 283.79 ± 72.43 | 342.29 ± 135.28 | 85.79 ± 25.56 | 81.49 ± 35.87 | **0.007** | 0.736 | 0.696 |
| T-CA | 944.20 ± 187.82 | 1697.34 ± 366.09 | 208.66 ± 50.32 | 459.47 ± 157.16^^ | **<0.001** | **0.030** | 0.265 |
| T-CDCA | 32.16 ± 5.80 | 38.92 ± 10.34 | 18.21 ± 4.32 | 16.67 ± 6.21 | **0.014** | 0.712 | 0.558 |
| T-DCA | 25.57 ± 4.31 | 48.23 ± 8.12* | 7.52 ± 1.87 | 16.26 ± 4.86^^^ | **<0.001** | **0.005** | 0.196 |
| T-UDCA | 15.87 ± 4.47 | 17.35 ± 5.96 | 6.68 ± 2.40 | 5.93 ± 2.99 | **0.019** | 0.931 | 0.790 |
| αMCA | 9.74 ± 6.68 | 3.56 ± 1.53 | 6.09 ± 4.88 | 3.10 ± 1.31 | 0.635 | 0.290 | 0.709 |
| βMCA | 42.66 ± 26.76 | 20.01 ± 7.08 | 12.99 ± 4.55 | 14.67 ± 4.12 | 0.225 | 0.464 | 0.397 |
| CA | 91.76 ± 80.61 | 79.71 ± 49.30 | 32.08 ± 26.44 | 26.24 ± 13.50 | 0.261 | 0.858 | 0.950 |
| CDCA | 3.54 ± 3.81 | 2.11 ± 1.86 | 7.67 ± 8.12 | 1.91 ± 1.24 | 0.674 | 0.442 | 0.643 |
| DCA | 12.59 ± 3.54 | 15.58 ± 2.05 | 7.31 ± 1.25 | 14.38 ± 2.77 | 0.212 | 0.056 | 0.431 |
| LCA | 8.50 ± 2.82 | 3.30 ± 0.94 | 3.17 ± 0.85 | 4.10 ± 0.63 | 0.160 | 0.185 | 0.059 |
| UDCA | 1.53 ± 1.10 | 1.06 ± 0.52 | 1.44 ± 0.53 | 1.34 ± 0.39 | 0.874 | 0.688 | 0.774 |
| G-TOTALS | 26.30 ± 8.36 | 23.11 ± 4.85 | 11.56 ± 3.73 | 11.58 ± 4.42 | **0.026** | 0.779 | 0.778 |
| T-TOTALS | 1363.92 ± 275.94 | 2223.43 ± 533.83 | 374.94 ± 90.89 | 616.86 ± 215.99 | **<0.001** | 0.096 | 0.345 |
| Unconj TOTALS | 170.31 ± 121.71 | 125.33 ± 60.55 | 70.76 ± 43.92 | 65.75 ± 20.38 | 0.277 | 0.731 | 0.783 |
| % G-TOTALS | 1.66 ± 0.36 | 1.53 ± 0.71 | 2.88 ± 1.10 | 2.46 ± 0.57 | 0.150 | 0.707 | 0.844 |
| % T-TOTALS | 87.36 ± 6.10 | 90.85 ± 3.20 | 82.72 ± 6.49 | 77.06 ± 6.93 | 0.125 | 0.854 | 0.441 |
| % Unconj TOTALS | 10.98 ± 5.85 | 7.62 ± 2.74 | 14.40 ± 5.99 | 20.48 ± 6.62 | 0.148 | 0.806 | 0.397 |
| 12-hydroxy/non-hydroxy | 2.38 ± 0.18 | 4.85 ± 0.53*** | 1.56 ± 0.23 | 3.04 ± 0.30*^^ | **<0.001** | **<0.001** | 0.153 |
| **TOTAL (ng/mL)** | 1560.53 ± 274.48 | 2371.87 ± 534.21 | 457.25 ± 93.92 | 694.18 ± 212.12^^ | **<0.001** | 0.112 | 0.378 |

Values are means ± SEM. Significance was determined by 2-way ANOVA (strain X diet) followed by Tukey's multiple comparisons test; *indicates effect of diet within strain (*p<0.05, **p<0.01, ***p<0.001); ^indicates effect of strain within diet (^p<0.05, ^^p<0.01, ^^^p<0.001).

**Supplemental Table S8.** Anthropometric, energy intake, and wheel running data from mice on a sedentary and volunteer wheel running mice.

| Variable | SED | | | VWR | | | P-value |
|---|---|---|---|---|---|---|---|
| Δ Body Mass (g) | 6.68 | ± | 0.75 | 3.86 | ± | 0.66 | **0.029** |
| Δ Fat Mass (g) | 6.81 | ± | 0.52 | 3.86 | ± | 0.62 | **0.022** |
| Δ Lean Mass (g) | -1.86 | ± | 0.37 | 0.35 | ± | 0.51 | **0.026** |
| Body Mass (g) | 36.31 | ± | 0.74 | 32.23 | ± | 1.02 | **0.001** |
| % Fat Mass | 0.28 | ± | 0.01 | 0.19 | ± | 0.02 | **0.039** |
| % Lean Mass | 0.64 | ± | 0.01 | 0.72 | ± | 0.02 | **0.006** |
| Avg Daily Energy Intake (kcal) | 11.78 | ± | 0.21 | 15.46 | ± | 0.42 | **0.001** |
| Avg Daily Running Distance (km) | - | | - | 10.02 | ± | 0.52 | |

Values are means ± SEM (n=7-8). Significance was determined by Ttest.

**Supplemental Table S9.** Male liver-specific Cyp7a1 KO anthropometric, energy intake, and wheel running data from sedentary and volunteer wheel running mice.

| | **Ctrl** | | **LCyp7a1KO** | | **P-Value** | | |
|---|---|---|---|---|---|---|---|
| | **SED** | **VWR** | **SED** | **VWR** | **Genotype** | **Exercise** | **Genotype x Exercise** |
| Body Weight (g) | 32.2 ± 2.2 | 32.3 ± 1.8 | 32.4 ± 1.4 | 34.6 ± 1.4 | 0.489 | 0.509 | 0.544 |
| Δ Body Weight (g) | 0.7 ± 0.4 | 0.8 ± 0.3 | -0.1 ± 0.4 | 0.4 ± 0.3 | 0.074 | 0.439 | 0.596 |
| Δ Fat Mass (g) | 0.6 ± 0.2 | -0.8 ± 0.9 | -1.0 ± 0.6 | -2.0 ± 0.9 | 0.061 | 0.114 | 0.773 |
| Δ Fat Free Mass (g) | -0.1 ± 0.3 | 0.3 ± 0.2 | 0.7 ± 0.4 | 1.2 ± 0.3 | **0.007** | 0.133 | 0.922 |
| Avg Daily Energy Intake (kcal) | 8.9 ± 0.4 | 11.7 ± 0.6 | 10.1 ± 0.4 | 12.5 ± 0.7 | 0.067 | **0.001** | 0.730 |
| Avg. Daily Running Distance (km) | - | 6.3 ± 1.3 | - | 5.6 ± 0.8 | 0.662 | - | - |

Values are means ± SEM (n=6-8). Significance was determined by 2-way ANOVA (Genotype X VWR) followed by Tukey's multiple comparisons test.

**Supplemental Table S10.** Female liver-specific Cyp7a1 KO anthropometric, energy intake, and wheel running data from sedentary and volunteer wheel running mice.

| | Ctrl | | LCyp7a1KO | | P-Value | | |
|---|---|---|---|---|---|---|---|
| | **SED** | **VWR** | **SED** | **VWR** | **Genotype** | **Exercise** | **Genotype x Exercise** |
| Body Weight (g) | 24.4 ± 1.8 | 22.7 ± 0.5 | 24.0 ± 1.3 | 23.5 ± 0.6 | 0.838 | 0.334 | 0.600 |
| Δ Body Weight (g) | 0.4 ± 0.3 | -0.4 ± 0.2 | 0.5 ± 0.2 | 0.5 ± 0.5 | 0.200 | 0.363 | 0.363 |
| Δ Fat Mass (g) | 0.2 ± 0.5 | -1.5 ± 0.9 | 0.5 ± 0.2 | -1.7 ± 0.5 | 0.983 | **0.008** | 0.722 |
| Δ Fat Free Mass (g) | 0.5 ± 0.2 | 0.9 ± 0.2 | -0.1 ± 0.3 | 0.3 ± 0.1 | **0.024** | 0.107 | 0.895 |
| Avg Daily Energy Intake (kcal) | 8.4 ± 0.5 | 10.3 ± 0.6 | 8.3 ± 0.4 | 11.0 ± 0.6 | 0.606 | **0.001** | 0.500 |
| Avg. Daily Running Distance (km) | - | 8.8 ± 0.9 | - | 8.6 ± 0.2 | 0.830 | - | - |

Values are means ± SEM (n=6-8). Significance was determined by 2-way ANOVA (Genotype X VWR) followed by Tukey's multiple comparisons test.

**Supplemental Table S11.** Male liver-specific Cyp7a1KO **l**iver bile acid content from sedentary and volunteer wheel-running mice.

| | **Ctrl** | | **LCyp7a1KO** | | **P-Value** | | |
|---|---|---|---|---|---|---|---|
| **(ug/g of liver)** | **SED** | **VWR** | **SED** | **VWR** | **Genotype** | **Exercise** | **Genotype x Exercise** |
| T-αMCA | 2.76 ± 0.46 | 1.20 ± 0.24* | 0.58 ± 0.15* | 0.32 ± 0.05[#] | **0.001** | **0.001** | **0.016** |
| T-βMCA | 11.11 ± 6.03 | 3.60 ± 0.65 | 4.02 ± 0.90 | 5.84 ± 0.66 | 0.370 | 0.293 | 0.091 |
| T-CA | 32.62 ± 11.53 | 16.28 ± 2.67 | 9.32 ± 2.34 | 11.17 ± 1.63 | **0.013** | 0.184 | 0.099 |
| T-CDCA | 3.39 ± 1.24 | 1.12 ± 0.18 | 0.60 ± 0.17 | 0.47 ± 0.13 | **0.004** | **0.039** | 0.062 |
| T-DCA | 3.26 ± 0.59 | 3.81 ± 0.54 | 1.05 ± 0.29 | 1.07 ± 0.18 | **0.001** | 0.505 | 0.527 |

Values are means ± SEM (n=6-8). Significance was determined by 2-way ANOVA (Genotype X VWR) followed by Tukey's multiple comparisons test.

**Supplemental Table S12.** Female liver-specific Cyp7a1KO liver bile acid content from sedentary and volunteer wheel-running mice.

| | **Ctrl** | | **LCyp7a1KO** | | **P-Value** | | |
|---|---|---|---|---|---|---|---|
| **(ug/g of liver)** | **SED** | **VWR** | **SED** | **VWR** | **Genotype** | **Exercise** | **Genotype x Exercise** |
| T-αMCA | 2.01 ± 0.41 | 2.19 ± 0.28 | 0.11 ± 0.03 | 0.88 ± 0.72 | **0.001** | 0.308 | 0.520 |
| T-βMCA | 5.51 ± 1.04 | 9.11 ± 1.62 | 3.77 ± 1.27 | 3.29 ± 0.64 | **0.005** | 0.216 | 0.110 |
| T-CA | 27.61 ± 5.29 | 48.25 ± 7.89 | 11.30 ± 2.70 | 16.58 ± 2.60 | **0.001** | **0.019** | 0.149 |
| T-CDCA | 2.55 ± 0.34 | 2.75 ± 0.40 | 0.21 ± 0.06 | 0.70 ± 0.41 | **0.001** | 0.314 | 0.668 |
| T-DCA | 4.31 ± 0.96 | 7.88 ± 1.64 | 0.36 ± 0.13 | 0.83 ± 0.20 | **0.001** | **0.048** | 0.124 |

Values are means ± SEM (n=6-8). Significance was determined by 2-way ANOVA (Genotype X VWR) followed by Tukey's multiple comparisons test.
