## Supplementary figures and images for "Aerobic capacity and exercise mediate protection against hepatic steatosis via enhanced bile acid metabolism"

### Supplemental Figure 1

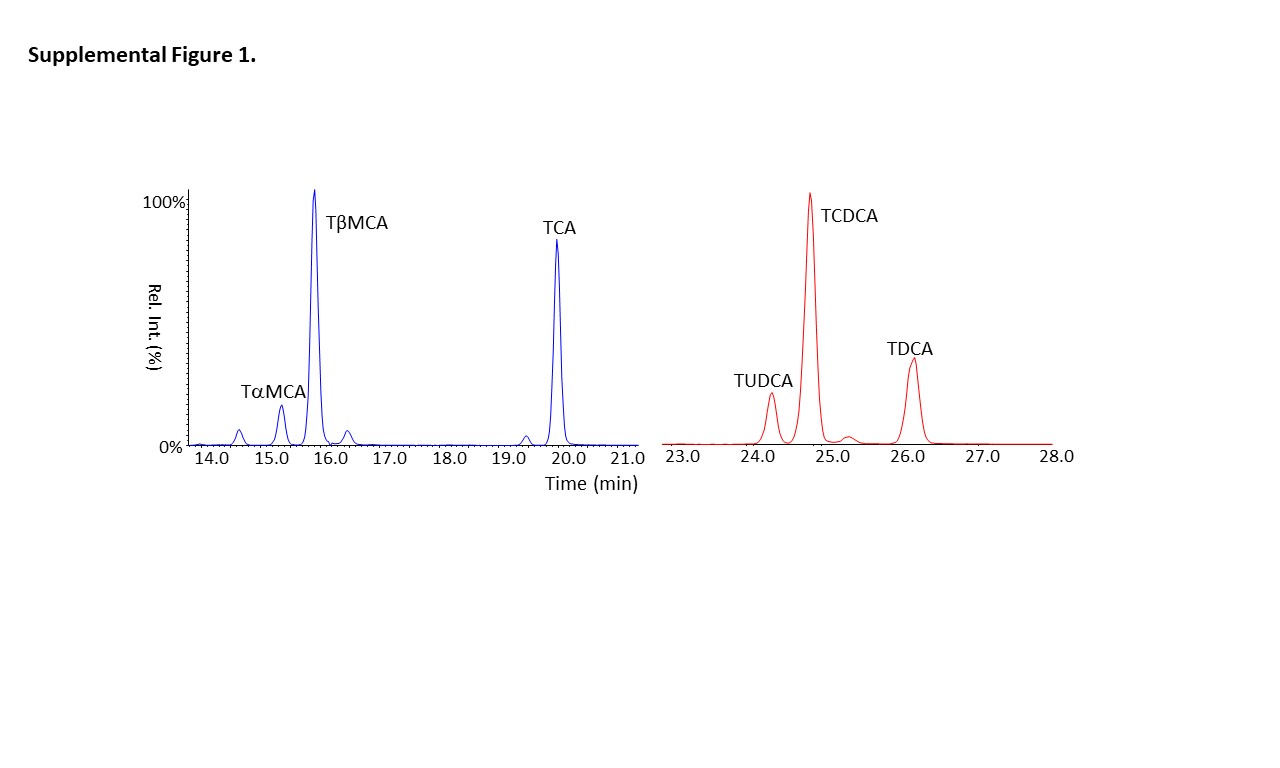

### Supplemental Figure 2

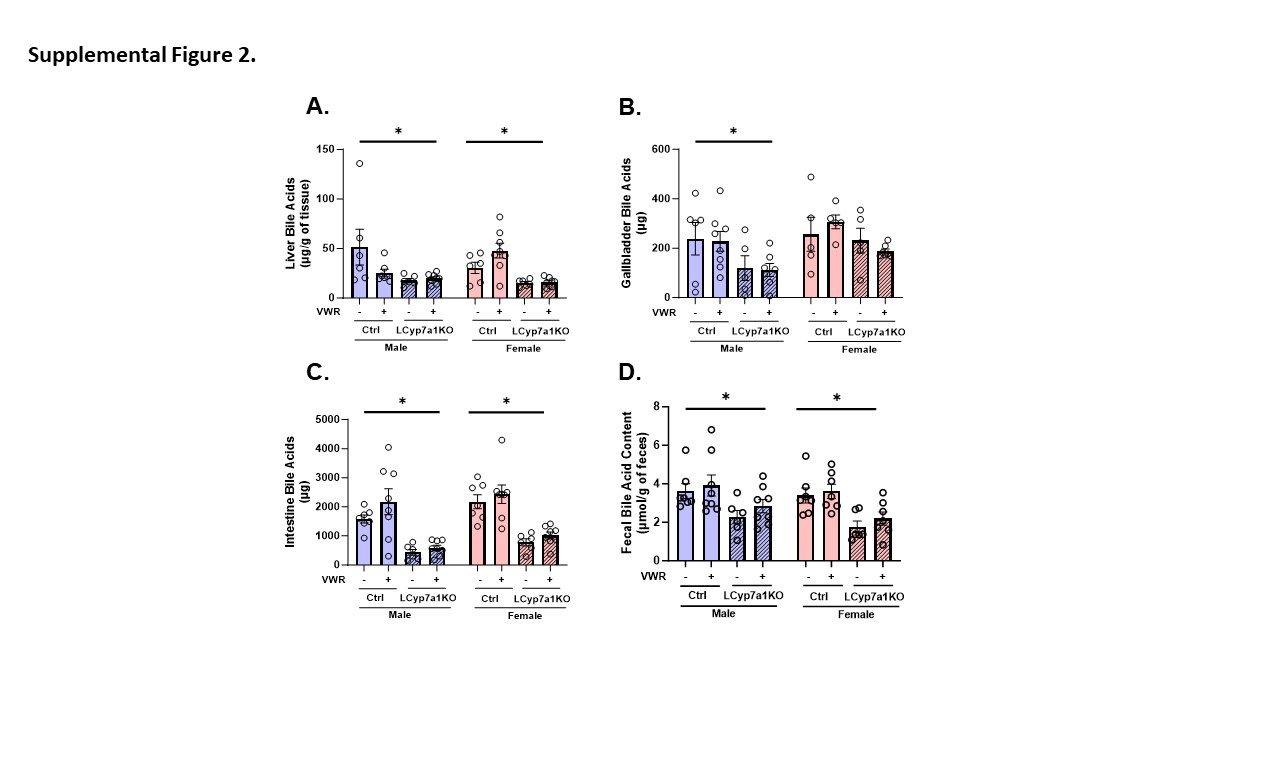
